# Dopaminergic Neurodegeneration Induced by Parkinson’s Disease-Linked G2019S LRRK2 is Dependent on Kinase and GTPase Activity

**DOI:** 10.1101/2019.12.17.879759

**Authors:** An Phu Tran Nguyen, Elpida Tsika, Kaela Kelly, Nathan Levine, Xi Chen, Andrew B. West, Sylviane Boularand, Pascal Barneoud, Darren J. Moore

**Author notes:** AC Immune SA, EPFL Innovation Park, 1015 Lausanne, Switzerland.

## Abstract

Mutations in the *leucine-rich repeat kinase 2* (*LRRK2*) gene cause late-onset, autosomal dominant familial Parkinson’s disease (PD) and represent the most common known cause of PD. LRRK2 can function as both a protein kinase and GTPase and PD-linked mutations are known to influence both of these enzymatic activities. While PD-linked LRRK2 mutations can commonly induce neuronal damage and toxicity in cellular models, the mechanisms underlying these pathogenic effects remain uncertain. Rodent models based upon familial LRRK2 mutations often lack the hallmark features of PD and robust neurodegenerative phenotypes in general. Here, we develop a robust pre-clinical model of PD in adult rats induced by the brain delivery of recombinant adenoviral vectors with neuronal-specific expression of full-length human LRRK2 harboring the most common G2019S mutation. In this model, G2019S LRRK2 induces the robust degeneration of substantia nigra dopaminergic neurons, a pathological hallmark of PD. Introduction of a stable kinase-inactive mutation or in-diet dosing with the selective kinase inhibitor, PF-360, attenuates neurodegeneration induced by G2019S LRRK2. Neuroprotection provided by pharmacological kinase inhibition is mediated by an unusual mechanism involving the selective and robust destabilization of human LRRK2 protein in the rat brain relative to endogenous LRRK2. Our study further demonstrates that dopaminergic neurodegeneration induced by G2019S LRRK2 critically requires normal GTPase activity. The introduction of hypothesis-testing mutations that increase GTP hydrolysis or impair GTP binding activity provide neuroprotection against G2019S LRRK2 via distinct mechanisms. Taken together, our data demonstrate that G2019S LRRK2 induces neurodegeneration *in vivo* via a mechanism that is dependent on kinase and GTPase activity. Our study provides a robust rodent model of *LRRK2*-linked PD and nominates kinase inhibition and modulation of GTPase activity as promising disease-modifying therapeutic targets.

## Introduction

Parkinson’s disease (PD) typically occurs in a sporadic manner through a complex interaction between genetic risk, aging and environmental exposure (1). A small proportion of PD cases are inherited and are known to be caused by mutations in at least 14 different genes (2). Among them, mutations in the *leucine-rich repeat kinase 2* (*LRRK2*) gene cause late-onset, autosomal dominant PD and represent the most common cause of familial PD (3–6). Genome-wide association studies further link common variants at the *LRRK2* locus with increased risk for sporadic PD (7, 8). LRRK2 has therefore emerged as an important player and therapeutic target for familial and sporadic PD. In mammals, LRRK2 is ubiquitously expressed with enrichment in kidney, lung and various peripheral immune cells, and is present in multiple cell types throughout the brain including neurons (9–11). LRRK2 is a large multi-domain protein containing two central enzymatic domains, a Ras-of-Complex (Roc) GTPase domain and a tyrosine kinase-like kinase domain, linked by a C-terminal-of-Roc (COR) domain and flanked by four protein interaction repeat domains (12). Familial PD-linked mutations cluster within the Roc-COR tandem (N1437H, R1441C/G/H, R1628P and Y1699C) and kinase (I2012T, G2019S and I2020T) domains of LRRK2 suggesting important roles for both enzymatic activities in the pathophysiology of PD.

LRRK2 can function as both a GTPase and kinase *in vitro* and in cells, with an intact GTPase domain and the capacity for GTP-binding being critically required for kinase activity (13–16). Familial PD-linked mutations in LRRK2 commonly increase its kinase activity in mammalian cells to varying degrees and promote substrate phosphorylation (i.e. a subset of Rab GTPases) and autophosphorylation (i.e. at Ser1292) (17–19). For the common G2019S mutation located within the kinase activation loop, the effect on kinase activity is direct, whereas Roc-COR domain mutations are considered to act indirectly by impairing GTP hydrolysis activity and thereby prolonging the GTP-bound ‘on’ state of LRRK2 (15, 20–24). While GTPase and kinase activities of LRRK2 are clearly altered by familial mutations, it is less clear whether or how these enzymatic activities contribute to neuronal toxicity induced by mutant LRRK2. For example, many studies have routinely employed kinase-inactive mutations at the kinase proton acceptor site (D1994) to block neuronal damage in primary culture models induced by PD-linked mutant LRRK2 (25–30). However, null mutations at the D1994 residue (i.e. D1994A/N/S) selectively destabilize LRRK2 protein in primary neurons versus cell lines (13, 26, 31, 32), thereby making these types of single-cell neuronal assay difficult to adequately control. Similarly, commonly used hypothesis-testing mutations that disrupt GDP/GTP-binding within the P-loop of the GTPase domain (K1347A or T1348N) tend to markedly impair LRRK2 protein stability in most cell types (13, 23). Due to this adverse impact on LRRK2 protein levels, the neuroprotective effects of genetically inhibiting kinase activity or GTP-binding in neuronal models based upon PD-linked LRRK2 mutants have been difficult to robustly demonstrate. Notwithstanding these concerns, it is generally accepted that neuronal toxicity in primary culture models induced by mutant LRRK2 is mediated via a kinase-dependent mechanism whereas the contribution of GTPase activity is less certain (12).

The molecular mechanisms underlying the pathogenic effects of familial LRRK2 mutations in the mammalian brain are poorly understood due largely to the lack of robust neurodegenerative phenotypes in most animal models (33). While certain transgenic mouse models with high-level overexpression of mutant LRRK2 can develop a modest yet late-onset loss of substantia nigra dopaminergic neurons (34–36), these models are generally the exception, with most LRRK2 transgenic or knockin models of PD developing only subtle if any neuropathology over their lifespan (12, 33). Viral-mediated gene transfer in the rodent brain using large-capacity vectors such as a herpes simplex virus (HSV) amplicon or human adenovirus serotype 5 (Ad5) has been successful in producing models with robust and progressive dopaminergic neurodegeneration induced by human G2019S LRRK2, occurring over a shorter more feasible timeframe i.e. 3-6 weeks (29, 32, 37). In the HSV-LRRK2 mouse model, G2019S LRRK2 induces dopaminergic neuronal loss that is kinase-dependent albeit based upon using the unstable D1994A mutation or non-selective LRRK2 kinase inhibitors (29). Similarly, the Ad5-LRRK2 rat model reveals neuropathology induced by G2019S LRRK2 in a kinase-dependent manner, again using the unstable D1994N mutation (32). While these prior studies tend to support an important role for LRRK2 kinase activity in mediating neurodegeneration in PD, there have been few studies in rodent models rigorously evaluating whether or how kinase or GTPase activity meaningfully contribute to neurodegeneration induced by familial PD-linked LRRK2 mutations.

Here, we extend our prior work and have optimized adenoviral production, titer and delivery site in the Ad5-LRRK2 rat model of PD to provide a robust genetic and pharmacological evaluation of the contribution of kinase, GTP-binding and GTP hydrolysis activities to nigrostriatal pathway dopaminergic neurodegeneration induced by the common G2019S mutation in LRRK2. Our study highlights the importance of kinase and GTPase activity of LRRK2 in mediating neurodegeneration *in vivo* and validates both enzymatic activities as key therapeutic targets for PD.

## Results

### Pilot study of G2019S LRRK2 kinase activity following the intranigral delivery of Ad5-LRRK2 vectors

To begin to evaluate the enzymatic mechanisms by which G2019S LRRK2 mediates neuronal cell death in the adult rat brain, we developed second generation E1, E2a, E3-deleted recombinant human Ad5 vectors expressing either eGFP or full-length 3xFLAG-tagged human LRRK2 variants from a neuronal-specific synapsin-1 promoter (32, 37). The neuronal-restricted transgene expression produced by this Ad5 vector permits an investigation of the cell-autonomous pathogenic effects of LRRK2 variants in nigrostriatal pathway dopaminergic neurons. The most common PD-linked G2019S mutation was employed either alone or in combination with two distinct kinase-inactive mutations, D1994N (proton acceptor site) or K1906M (ATP-binding site), located in the kinase domain of LRRK2 (12). Prior studies by us and others have revealed that null mutations at the D1994 residue destabilize the LRRK2 protein selectively in primary neurons and brain tissue, whereas mutations at the K1906 residue are generally well-tolerated and stable (13, 31, 32). Therefore, we elected to compare the effects of these distinct kinase-inactive mutations on G2019S LRRK2-induced phenotypes. Purified Ad5 vectors were able to infect human SH-SY5Y neural cells in a titer-dependent manner and, most importantly, produce equivalent levels of each FLAG-LRRK2 variant (G2019S, G2019S/D1994N and G2019S/K1906M), as shown by Western blot analysis (Fig. 1A). We next conducted pilot studies in the adult rat brain by the stereotactic unilateral delivery of these Ad5 vectors to the substantia nigra pars compacta (SNpc) using a maximal viral titer of ∼4.5 x 10^9^ viral particles (vp), similar to our previous studies (32, 37). We opted to evaluate direct delivery of Ad5 vectors at a single site in the SNpc, since our prior study using a similar viral titer produced inefficient retrograde axonal transport of viral particles from the striatum to the SNpc (32). At 21 days post-injection, numerous neurons labeled with GFP or LRRK2 variants (FLAG) are detected within the ipsilateral nigra largely confined to the region spanning ± 80 μm from the injection site, suggesting limited viral transduction throughout the entire substantia nigra at this titer (Fig. 1B). Consistent with our prior studies (13, 32), we detect qualitatively equivalent levels of G2019S and G2019S/K1906M LRRK2 proteins whereas G2019S/D1994N LRRK2 protein is detected at markedly lower levels, indicating the reduced stability of this variant in the brain (Fig. 1B). To evaluate the impact of these LRRK2 variants on neurodegeneration, unbiased stereology was used to estimate the number of tyrosine hydroxylase (TH)-positive dopaminergic neurons in the ipsilateral versus contralateral SNpc at 21 days after Ad5 delivery (Fig. 1C). G2019S LRRK2 induces a marked significant loss of nigral dopaminergic neurons (42.7 ± 12.9% loss) compared to an eGFP control (20.7 ± 12.9% loss). Notably, G2019S/D1994N and G2019S/K1906M LRRK2 both attenuate neuronal loss compared to G2019S LRRK2, producing levels comparable to eGFP (25.9 ± 14.9 and 23.8 ± 10.7% loss, respectively), although these reductions do not reach significance due to the limited number of animals used (Fig. 1D). Together, these pilot data validate the intranigral delivery of Ad5 vectors as a potential approach for modeling dopaminergic neurodegeneration induced by G2019S LRRK2, and initially suggest that these pathogenic effects are kinase-dependent based upon the stable kinase-inactive K1906M mutation.

**Figure 1.**
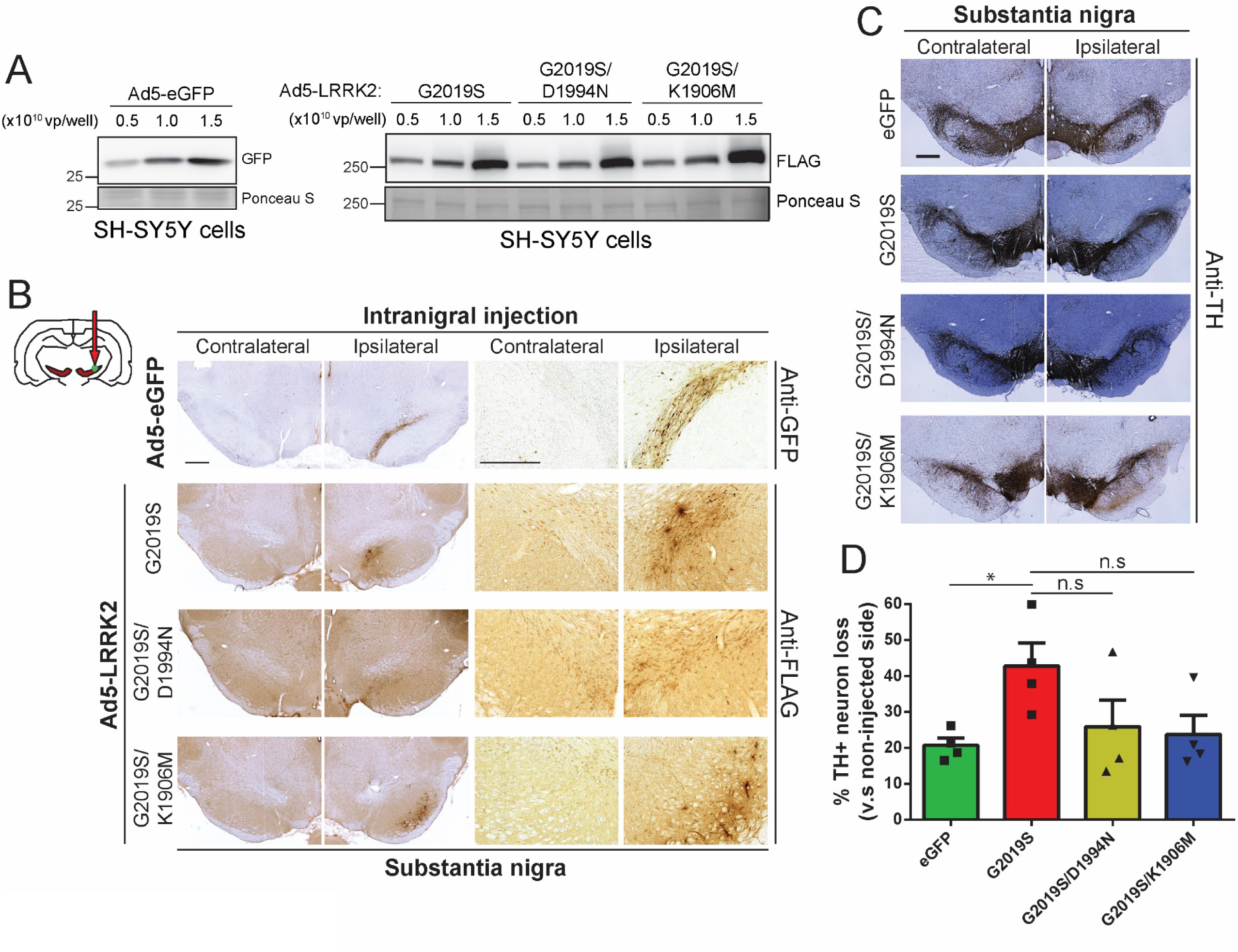
Pilot studies evaluating role of kinase activity using new Ad5-LRRK2-G2019S vectors. **A)** Validation of Ad5 vectors in SH-SY5Y cells. Cells were transduced with increasing viral particles (vp) of each Ad5-eGFP or Ad5-LRRK2 vector (0.5, 1 or 1.5 x 10^10^ vp/well). Western blot analysis of cell extracts with GFP or FLAG antibody to detect a titer-dependent increase in human LRRK2 levels. Ponceau S stain was used as a protein loading control. Molecular mass markers are indicated in kilodaltons. **B)** Ad5-eGFP and Ad5-LRRK2 (G2019S, G2019S/D1994N or G2019S/K1906M) vectors were delivered at a single injection site (4.5 x 10^9^ vp/site, in 2.5 µl) in the ipsilateral substantia nigra pars compacta of rats, as indicated (red arrow). Immunohistochemistry for eGFP (anti-GFP antibody) or human LRRK2 variants (anti-FLAG antibody) in the ipsilateral substantia nigra at 10 days post-injection. Images indicate low and high magnification for each vector. Scale bar: 500 μm. **C)** Immunohistochemistry revealing dopaminergic neurons (anti-TH antibody) in contralateral and ipsilateral substantia nigra at 21 days post-delivery of Ad5 vectors. All sections were counterstained with Cresyl violet. **D)** Analysis of neurodegeneration in rat substantia nigra at 21 days. TH-positive dopaminergic neuron number in the substantia nigra assessed by unbiased stereology. Bars represent % neuronal loss in the injected ipsilateral nigra relative to the contralateral nigra (mean ± SEM, *n* = 4 animals/vector), \**P*<0.05 by one-way ANOVA with Bonferroni’s multiple comparisons test. *n.s*, non-significant.

### Optimization of Ad5-LRRK2 vector titer and route of delivery in the rat brain

We next sought to optimize and improve Ad5 vector production, titer and delivery to further enhance the expression levels of human LRRK2 variants (WT, G2019S and G2019S/K1906M) in nigral dopaminergic neurons of this rat model. We elected to move ahead with the kinase-inactive K1906M mutant due to its improved protein stability in the brain relative to the D1994N mutant. We decided to compare two Ad5 delivery paradigms (intranigral versus intrastriatal) since both approaches have proven successful in producing human LRRK2 expression in nigral dopaminergic neurons at a viral titer of ∼4.5 x 10^9^ vp/site (Fig. 1 and (37)), albeit with some limitations in transduction efficiency. For example, the limited number of nigral neurons locally transduced following intranigral Ad5 delivery may relate to the low somatic expression in dopaminergic neurons of the coxsackie and adenovirus receptors (CARs) that are critical for efficient viral uptake (38). Alternatively, intrastriatal Ad5 delivery can produce inefficient retrograde transport of viral particles along dopaminergic axons to the SNpc, subsequently minimizing G2019S LRRK2-induced neurodegeneration (32). To overcome these issues, we successfully improved the titer of Ad5 vectors by optimizing different steps in the viral amplification and purification procedures. These improvements lead to a 3-4-fold increase in the stock titer of Ad5-LRRK2 vectors (WT, G2019S, G2019S/K1906M) to 6-8 x 10^9^ vp/μl (from ∼2 x 10^9^ vp/µl), which would allow the maximal delivery of up to ∼1.6 x 10^10^ vp/site (in a 2 μl volume) in both injection paradigms. To first evaluate these improved Ad5-LRRK2 preparations, we demonstrate the titer-dependent and equivalent expression levels of G2019S and G2019S/K1906M FLAG-LRRK2 variants in SH-SY5Y cells by Western blot analysis (Fig. 2A) whereas WT LRRK2 levels are modestly lower. We confirm the markedly increased kinase activity of G2019S LRRK2 relative to WT LRRK2 in cells by monitoring autophosphorylation at Ser1292, whereas G2019S/K1906M LRRK2 is kinase-inactive (Fig. 2A). Constitutive phosphorylation at Ser910 and Ser935 is not altered between these LRRK2 kinase-active and kinase-inactive variants (Fig. 2A), consistent with prior reports (39). Our data confirm the kinase activity status (pSer1292) of our Ad5-LRRK2 vectors indicating that G2019S is hyperactive and G2019S/K1906M is inactive.

**Figure 2.**
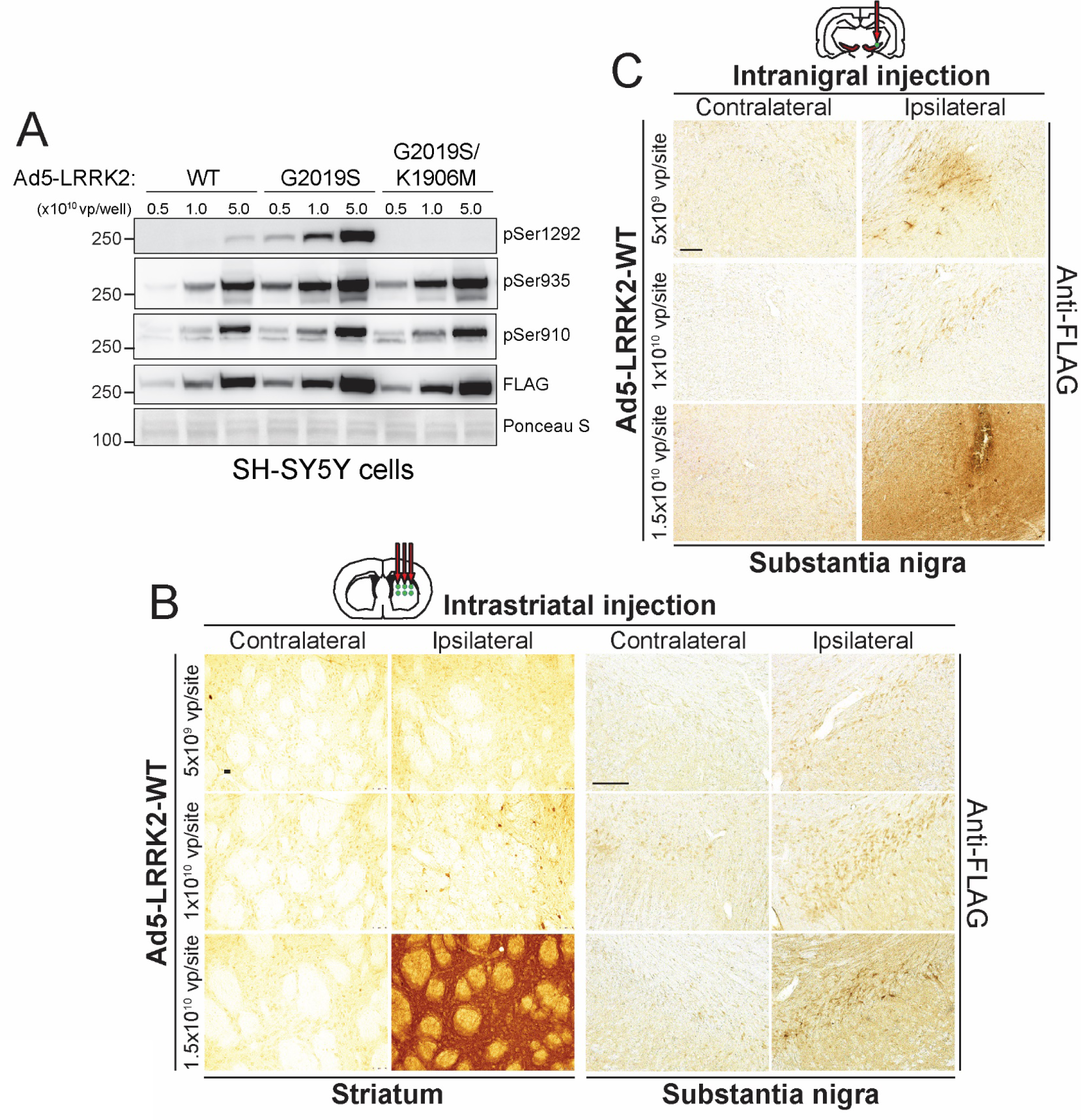
Evaluation of Ad5 vector infectivity in rat brain via distinct delivery paradigms. **A)** Validation of new high-titer Ad5-LRRK2 (WT, G2019S and G2019S/K1906M) vectors and LRRK2 phosphorylation in SH-SY5Y cells. Cells were transduced with increasing viral particles (vp) of each Ad5-LRRK2 vector (0.5, 1 or 5.0 x 10^10^ vp/well). Western blot analysis of cell extracts with FLAG antibody (transduced human LRRK2) or antibodies to detect LRRK2 autophosphorylation (pSer1292) or constitutive phosphorylation (pSer935, pSer910) sites to detect a titer-dependent increase in human LRRK2 levels/activity. Ponceau S stain was used as a protein loading control. Molecular mass markers are indicated in kilodaltons. **B)** Evaluation of Ad5-LRRK2 vectors following intrastriatal delivery with increasing viral titer (0.5, 1 or 1.5 x 10^10^ vp/site, at 2.5 μl/site) at six distinct injection sites. Immunohistochemistry showing dose-dependent expression of human LRRK2-WT (anti-FLAG antibody) in the ipsilateral rat striatum and substantia nigra at 10 days post-injection. Scale bars: 500 μm. **C)** Evaluation of Ad5-LRRK2 vectors following intranigral delivery with increasing viral titer (0.5, 1 or 1.5 x 10^10^ vp/site, in 2.5 μl) at a single injection site. Immunohistochemistry showing dose-dependent expression of human LRRK2-WT (anti-FLAG antibody) in rat substantia nigra at 10 days post-injection. Scale bars: 500 μm.

To evaluate the transduction efficiency of these new Ad5-LRRK2 preparations *in vivo*, we injected adult rats using intrastriatal (6 sites, 2 μl/site) or intranigral (single site, 2 μl) delivery paradigms with an Ad5-LRRK2 WT vector at three different titers (∼5 x 10^9^, 1 x 10^10^ and 1.5 x 10^10^ vp/site) (Fig. 2B-C). At 10 days post-injection, we observe a titer-dependent expression of FLAG-LRRK2 in neurons and neuritic processes of the ipsilateral striatum and SNpc following intrastriatal Ad5 delivery (Fig. 2B) and increased labeling of FLAG-LRRK2 in the ipsilateral SNpc following intranigral delivery (Fig. 2C). LRRK2-positive neurons or neurites are not generally detected in the contralateral striatum or SNpc using either delivery method or at any viral titer (Fig. 2B-C). While FLAG-LRRK2-positive nigral dopaminergic neurons can already be detected following intrastriatal delivery of Ad5 at the lowest titer of 5 x 10^9^ vp/site, we observe a titer-dependent increase of LRRK2-positive nigral neuron number, suggesting a marked improvement in retrograde axonal transport of viral particles with increasing amounts of Ad5 (Fig. 2B). With intranigral delivery, using Ad5 at 5 x 10^9^ and 1 x 10^10^ vp/site results in LRRK2-positive expression that is limited around the injection site in the SNpc (Fig. 2C). However, a substantially greater area of the SNpc containing LRRK2-positive neuronal soma and neurites is transduced by Ad5 at the highest titer of 1.5 x 10^10^ vp/site, suggesting a broader diffusion of viral particles at this dose (Fig. 2C). Intrastriatal delivery produces abundant LRRK2-positive labeling limited to nigral dopaminergic neuronal soma throughout the entire rostro-caudal axis of the ipsilateral SNpc whereas intranigral delivery displays a more limited distribution to neurons and neurites within the SNpc (Fig. 2B-C). Using a similar approach, we confirm the selective transduction of nigral dopaminergic neurons by confocal co-localization following delivery of a new high-titer Ad5-eGFP vector to the rat striatum at the maximal viral titer of ∼1.5 x 10^10^ vp/site (Fig. S1). Taken together, our pilot studies varying titer and delivery site of Ad5 vectors demonstrate that intranigral and intrastriatal delivery routes in the adult rat brain are sufficient to achieve LRRK2 expression in nigral dopaminergic neurons in a viral titer-dependent manner. Accordingly, we have elected to use the maximal Ad5 titer of 1.5 x 10^10^ vp/site for all subsequent studies to obtain the highest expression of human LRRK2 throughout the nigrostriatal pathway.

### Analysis of protein levels and kinase activity of Ad5-LRRK2 variants in the rat brain

Prior to exploring the role of kinase activity in driving neurodegeneration, we first sought to confirm the protein levels and kinase activity status of the Ad5-LRRK2 vectors in the rat brain. To obtain the highest expression of human LRRK2 variants, we employed the intrastriatal delivery of Ad5-LRRK2 vectors (WT, G2019S and G2019S/K1906M) in adult rats (Fig. S2). At 10 days post-injection, the ipsilateral striatum was subjected to sequential detergent extraction to produce 1% Triton-soluble and Triton-insoluble (2% SDS-soluble) fractions. Human LRRK2 variants are detected in both Triton-soluble and insoluble fractions where the steady-state levels of G2019S and G2019S/K1906M LRRK2 proteins are equivalent by Western blot analysis whereas the level of WT LRRK2 protein is modestly lower (Fig. S2A). These data are similar to our observations in SH-SY5Y cells (Fig. 2A), indicating that genetic inactivation of kinase activity by the K1906M mutation does not alter the protein stability of G2019S LRRK2 in neural tissue and is generally well-tolerated.

We next sought to monitor LRRK2 kinase activity in brain extracts by evaluating Rab10 phosphorylation (pThr73-Rab10), a well-established substrate of LRRK2 (18, 40). While endogenous Rab10 is readily detected in Triton-soluble and insoluble fractions from striatum, we detect minimal levels of pT73-Rab10 that are not altered between extracts expressing WT, G2019S or G2019S/K1906M LRRK2 (Fig. S2A). These data suggest that pT73-Rab10 does not serve as a sensitive readout of LRRK2 kinase activity in the brain of this rat model, similar to prior reports in rodent brain (41). Instead, LRRK2 kinase activity was monitored by autophosphorylation at Ser1292 in brain tissue. Following the immunoprecipitation of FLAG-LRRK2 variants from soluble striatal extracts, we detect a ∼2-fold increase of pSer1292 levels for G2019S LRRK2 compared to WT LRRK2 by Western blot analysis, and a near complete absence of pSer1292 signal for the G2019S/K1906M LRRK2 protein (Fig. S2B). The corresponding levels of constitutive Ser935 phosphorylation are not altered between these LRRK2 variants (Fig. S2B) as expected (39). These findings confirm our observations in SH-SY5Y cells (Fig. 2A) and indicate that G2019S LRRK2 is kinase-hyperactive whereas G2019S/K1906M LRRK2 is kinase-inactive in brain tissue. We next attempted to validate the alterations in pSer1292-LRRK2 levels by immunofluorescent labeling and confocal microscopy in the striatum and substantia nigra following intrastriatal delivery of Ad5-LRRK2 vectors (Fig. S2C). At 42 days post-injection, we can readily detect small numbers of FLAG-LRRK2-positive neurons in the ipsilateral striatum and SNpc yet these neurons exhibit minimal labeling for pSer1292-LRRK2. Moreover, the levels of pSer1292-LRRK2 do not differ between neurons expressing G2019S or G2019S/K1906M LRRK2 (Fig. S2C) suggesting that the pS1292-LRRK2 antibody is not sufficiently sensitive for immunofluorescent detection in this rat model. Overall, our data provide evidence for increased kinase activity of G2019S LRRK2 in the rat brain that is inhibited by addition of the K1906M mutation.

### G2019S LRRK2 induces dopaminergic neurodegeneration in rats via a kinase-dependent mechanism

To extend our pilot data (Fig. 1) and rigorously evaluate the contribution of LRRK2 kinase activity in driving dopaminergic neurodegeneration, we employed our optimized delivery paradigms (intranigral versus intrastriatal, Fig. 2) and increased animal cohort sizes to compare between Ad5-LRRK2 variants (WT, G2019S and G2019S/K1906M). Adult rats were first subjected to a single intranigral injection of Ad5 vectors (∼1.5 x 10^10^ vp) and assessed for neurodegeneration after 21 days (Fig. 3). We observe the robust labeling of neurons and neurites with FLAG-LRRK2 variants or eGFP in the SNpc (Fig. 3A) comparable to the levels observed at 10 days post-injection (Fig. 2C). We find equivalent levels of FLAG-LRRK2-positive labeling between animals expressing the G2019S and G2019S/K1906M LRRK2 proteins (Fig. 3A), further confirming the normal protein stability of the G2019S/K1906M variant in brain tissue. Unbiased stereological counting reveals robust dopaminergic neurodegeneration induced by G2019S LRRK2 (36.9 ± 15.6% loss) in the ipsilateral SNpc that is only modestly reduced with WT LRRK2 (28.4 ± 17.0% loss), whereas neuronal loss is significantly attenuated with G2019S/K1906M LRRK2 (16.0 ± 12.4% loss) (Fig. 3B-C). The corresponding loss of total Nissl-positive neurons in the ipsilateral SNpc for each LRRK2 variant, albeit non-significant, suggests that LRRK2 variants induce neuronal cell death rather than altering TH phenotype (Fig. 3C), as previously reported (37). While there is a significant positive correlation between TH-positive and Nissl-positive neuronal loss in general within these animal cohorts (R^2^ = 0.2331, *P*<0.05), for unknown reasons this relationship does not always correlate for individual animals (Fig. 3D). Our data provide further evidence that G2019S LRRK2 induces dopaminergic neurodegeneration in rats via a mechanism that requires kinase activity following the direct intranigral delivery of Ad5 vectors. However, at this maximal Ad5 titer, there is already a high baseline of neuronal loss and differences between LRRK2 variants are rather modest.

**Figure 3.**
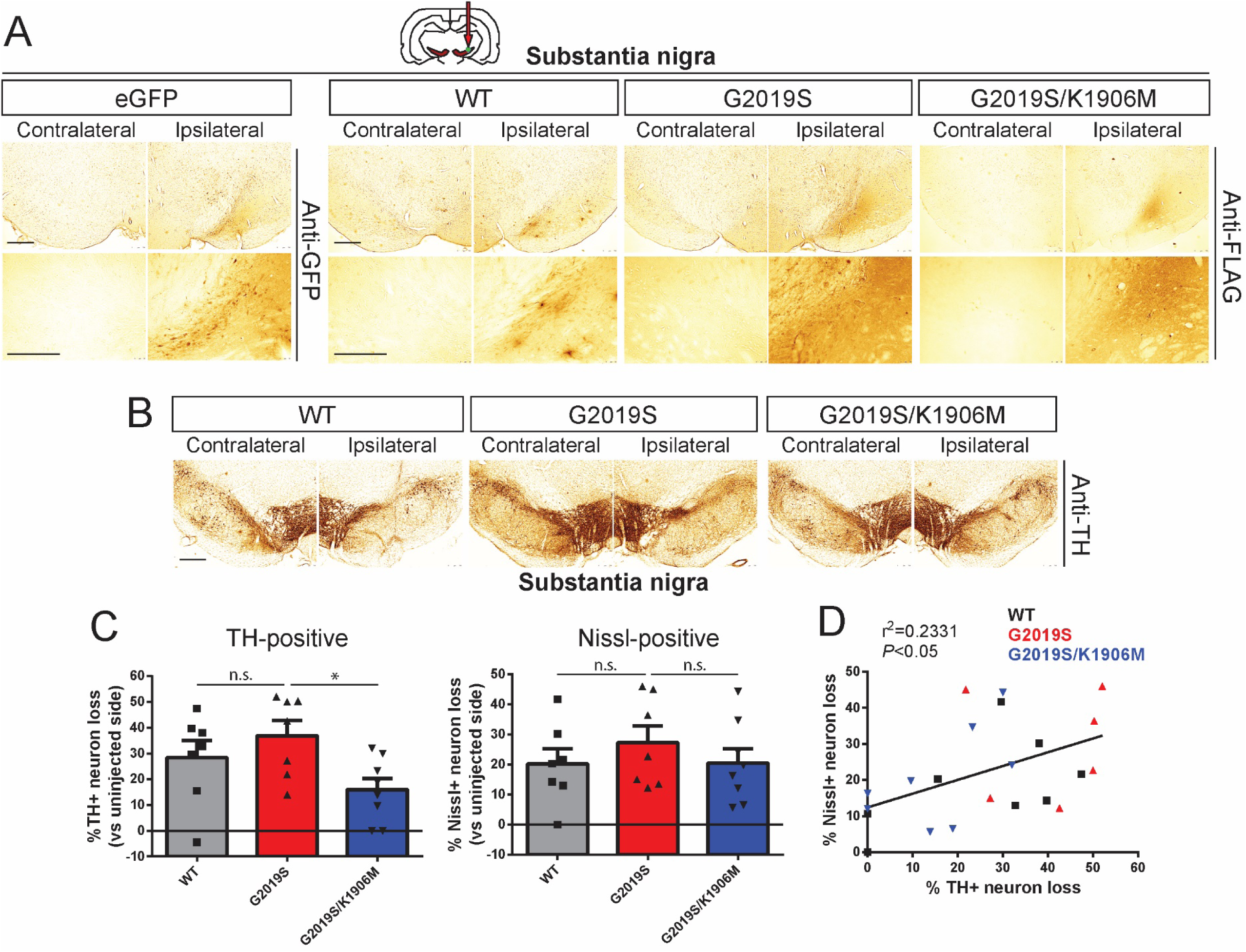
Intranigral delivery of Ad5-LRRK2-G2019S induces dopaminergic neurodegeneration in a kinase-dependent manner in adult rat brain. **A)** Ad5-LRRK2 (WT, G2019S or G2019S/K1906M) vectors were unilaterally delivered at a single site (1.5 x 10^10^ vp/site, in 2.5 μl) in the rat substantia nigra. Immunohistochemistry reveals expression of eGFP (anti-GFP antibody) or human LRRK2 variants (anti-FLAG antibody) that persists up to 21 days post-injection. Images indicate low and high magnification for each vector. Scale bars: 500 μm. **B)** Immunohistochemistry revealing dopaminergic neurons (anti-TH antibody) in contralateral and ipsilateral substantia nigra at 21 days post-delivery of Ad5 vectors. Scale bars: 500 μm. **C)** Unbiased stereological analysis of TH-positive dopaminergic and total Nissl-positive neuron number in the substantia nigra at 21 days post-injection. Bars represent % neuronal loss in the injected ipsilateral nigra relative to the contralateral nigra (mean ± SEM, *n* = 7-8 animals/vector), \**P*<0.05 by one-way ANOVA with Bonferroni’s multiple comparisons test. *n.s*, non-significant. **D)** Linear regression analysis indicating a significant positive correlation between TH-positive and Nissl-positive neuron loss in the substantia nigra. Data points represent individual animals. Pearson’s correlation coefficient, r^2^ = 0.2331, *P*<0.05.

We next conducted similar studies using intrastriatal delivery of Ad5-LRRK2 vectors (∼1.5 x 10^10^ vp/site at 6 distinct sites) in adult rats and evaluated neurodegeneration at 42 days post-injection (Fig. 4). Using a hexon protein-specific antibody that detects the capsid protein of Ad5 particles, we confirm equivalent amounts of each Ad5-LRRK2 vector following delivery to the ipsilateral striatum (Fig. S3A). FLAG-LRRK2 variants or eGFP are robustly detected in neurons and neurites of the ipsilateral striatum and within dopaminergic neurons of the ipsilateral SNpc at 42 days post-injection, with equivalent levels of G2019S and G2019S/K1906M LRRK2 proteins (Fig. 4A). FLAG-LRRK2-positive staining in the ipsilateral striatum and SNpc at 42 days is qualitatively reduced compared to pilot studies at 10 days post-injection (Fig. 2B) indicating that the pool of adenoviral particles undergoes clearance over this extended period. Unbiased stereology confirms robust nigral dopaminergic neurodegeneration in this model induced by G2019S LRRK2 (41.7 ± 14.8% neuronal loss) that is significantly and comparably attenuated with WT LRRK2 (17.1 ± 19.1%) or G2019S/K1906M LRRK2 (15.8 ± 20.1%) (Fig. 4B-C). TH-positive neuronal loss induced by each LRRK2 variant is supported by a parallel loss of total Nissl-positive neurons in the ipsilateral SNpc (Fig. 4C), with a significant positive correlation between these neuronal counts within these animal cohorts (R^2^ = 0.3433, *P*<0.05; Fig. 4D). Our data further demonstrate a key requirement of kinase activity for the robust dopaminergic neurodegeneration induced by G2019S LRRK2 in the rat brain following the intrastriatal delivery of Ad5 vectors. Although the extent of neuronal loss induced by G2019S LRRK2 is comparable between intranigral and intrastriatal Ad5 delivery paradigms, a key difference lies in the more modest impact of WT LRRK2 in the intrastriatal model whereas G2019S/K1906M LRRK2 exhibits a consistently low level of neuronal loss in both delivery paradigms. The delivery of Ad5-eGFP vectors at this maximal viral titer produces extensive non-specific neuronal toxicity in the substantia nigra, a problem commonly encountered in other viral-based rodent models (42), indicating that GFP does not serve as a useful negative control *in vivo*. Instead, we have used kinase-inactive G2019S/K1906M LRRK2 to set our baseline levels of neuronal toxicity in this rat model to control for the non-specific effects due to LRRK2 overexpression and/or Ad5 infection (Fig. 3-4). Taken together, these data indicate that the intrastriatal delivery of Ad5-LRRK2 vectors produces a more robust rodent model of neurodegeneration in which to identify a critical contribution of kinase activity.

**Figure 4.**
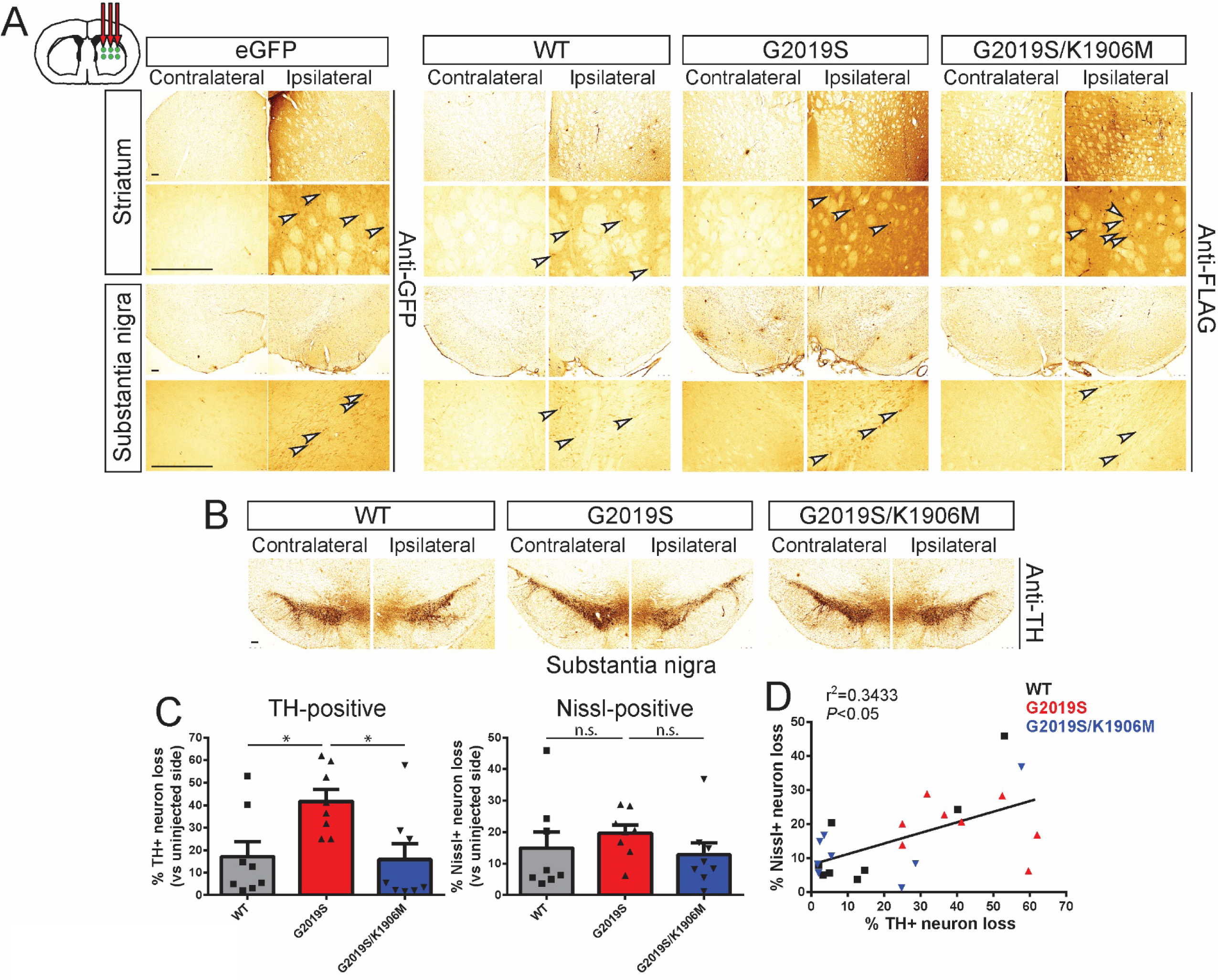
Intrastriatal delivery of Ad5-LRRK2-G2019S induces kinase-dependent dopaminergic neurodegeneration in adult rat brain. **A)** Ad5-LRRK2 (WT, G2019S and G2019S/K1906M) vectors were unilaterally delivered at six injection sites (1.5 x 10^10^ vp/site, in 2.5 μl) to the rat striatum. Immunohistochemistry reveals expression of eGFP (anti-GFP antibody) or human LRRK2 variants (anti-FLAG antibody) in the ipsilateral striatum and nigra that persists up to 42 days post-injection. Images indicate low and high magnification for each vector and brain region. Scale bars: 500 μm. **B)** Immunohistochemistry revealing dopaminergic neurons (anti-TH antibody) in contralateral and ipsilateral substantia nigra at 42 days post-delivery of Ad5 vectors. All sections were counterstained with Cresyl violet. Scale bars: 500 μm. **C)** Unbiased stereological analysis of TH-positive dopaminergic and total Nissl-positive neuron number in the substantia nigra at 42 days post-injection. Bars represent % neuronal loss in the injected ipsilateral nigra relative to the contralateral nigra (mean ± SEM, *n* = 8 animals/vector), \**P*<0.05 by one-way ANOVA with Bonferroni’s multiple comparisons test. *n.s*, non-significant. **D)** Linear regression analysis indicating a significant positive correlation between TH-positive and Nissl-positive neuron loss in the substantia nigra. Data points represent individual animals. Pearson’s correlation coefficient, r^2^ = 0.3433, *P*<0.05.

### Evaluation of PD-relevant neuropathological markers in the Ad5-LRRK2 rat model

Our prior study using intrastriatal delivery of Ad5-LRRK2 vectors at a lower viral titer (∼4.5 x 10^9^ vp/site) demonstrated that G2019S LRRK2 could induce striatal pathology in a kinase-dependent manner, including neurite degeneration (Gallyas silver stain), ubiquitin-positive neuronal inclusions and abnormal phospho-neurofilament distribution (32). To evaluate the extent of neuropathology throughout the nigrostriatal pathway induced by Ad5-LRRK2 vector delivery at a 3-fold higher titer (∼1.5 x 10^10^ vp/site) in this study, we assessed a number of markers to monitor for protein inclusions or aggregates, neurite and axonal damage, and microglial activation (Fig. S3-S6). In rat brains subjected to intrastriatal delivery of Ad5-LRRK2 vectors, we observe an increase in ubiquitin immunoreactivity in the ipsilateral striatum at 42 days although this is equivalent between WT, G2019S and G2019S/K1906M LRRK2 suggesting it arises in a manner independent of LRRK2 or its kinase activity (Fig. S3B). In contrast, G2019S LRRK2 specifically increases APP-positive inclusions in the ipsilateral striatum, a sensitive marker of axonal damage (43, 44), that is absent with WT or G2019S/K1906M LRRK2 variants (Fig. S3C). G2019S LRRK2 also markedly increases silver-positive neurites in the striatum, a marker of neurite degeneration (44–46), relative to WT or G2019S/K1906M LRRK2 (Fig. S3D). These data indicate that G2019S LRRK2 induces axonal degeneration via a kinase-dependent mechanism in this rat model, consistent with our prior study (32). This neuropathology was generally absent from the ipsilateral SNpc suggesting that it is confined to the striatum where human LRRK2 levels are highest (data not shown). Consistent with our prior study (32), α-synuclein pathology is not induced by G2019S LRRK2 in this rat model following the evaluation of α-synuclein levels, Ser129 phosphorylation or insoluble species in the striatum and ventral midbrain compared to WT or G2019S/K1906M LRRK2 (Fig. S4). While we do observe increased immunoreactivity for phosphorylated tau (pSer202/pThr205; AT8) in the ipsilateral SNpc, this is most likely non-specific, as it occurs equivalently with all LRRK2 variants and does not result in alterations in tau levels, phosphorylation or detergent solubility in extracts from striatum and ventral midbrain (Fig. S5). Microglial activation is detected in the ipsilateral striatum and SNpc as indicated by the modest increase of Iba1-positive immunoreactivity induced equivalently by all Ad5-LRRK2 vectors at 42 days following their intrastriatal delivery (Fig. S6A-B). In comparison, robust microglial activation is evident in the SNpc at 21 days following the intranigral delivery of Ad5-LRRK2 vectors (Fig. S6C), indicating a more pronounced inflammatory response to Ad5 delivery in the SNpc relative to the striatum. Microglial activation occurs independent of LRRK2 kinase activity in the rat brain and is most likely due to Ad5 infection. Collectively, these data demonstrate that G2019S LRRK2 selectively induces modest axonal pathology in the striatum via a kinase-dependent mechanism, whereas ubiquitin accumulation, tau hyperphosphorylation and microglial activation in this model are unrelated to LRRK2 kinase activity.

### Pharmacological inhibition of kinase activity with PF-360 protects against G2019S LRRK2 in rats

To further evaluate the contribution of kinase activity to the pathogenic effects of G2019S LRRK2 in this rat model, we explored the neuroprotective effects of a third generation, selective, potent and brain-permeable LRRK2 kinase inhibitor, PF-06685360 (PF-360) (47). Rodent chow was developed containing PF-360 at either 35 or 175 mg per kg of chow, as recently described (41), that was previously shown to be well-tolerated and produce complete inhibition of LRRK2 kinase activity in lung and near-complete inhibition in brain of rats. Rats were first subjected to the unilateral intrastriatal delivery of Ad5-LRRK2 G2019S vector followed by in-diet administration of PF-360 for 7 days beginning at 7 days post-injection (Fig. 5A). PF-360 administration at 35 mg/kg (equivalent to ∼3.2 mg/kg/day) produces a significant yet modest inhibition of endogenous LRRK2 kinase activity in brain by monitoring pS935-LRRK2 levels in detergent extracts from the contralateral striatum with negligible effects on total LRRK2 levels (Fig. 5B). In contrast, human G2019S LRRK2 levels are dramatically reduced in the ipsilateral striatum by PF-360 at this low dose (Fig. 5B). With PF-360 dosing at 175 mg/kg (equivalent to ∼16 mg/kg/day), endogenous pS935-LRRK2 is almost fully inhibited in the brain yet total LRRK2 levels remain unaffected, whereas human G2019S LRRK2 levels are further reduced (Fig. 5C). These data suggest that human G2019S LRRK2 is selectively and markedly destabilized in the rat brain by PF-360 administration relative to endogenous LRRK2. Administration of PF-360 (175 mg/kg chow) also results in near maximal levels of LRRK2 kinase inhibition (pS935-LRRK2) in peripheral tissues including lung, kidney, spleen and PBMCs yet with minimal effects on the total levels of endogenous LRRK2 at this dose (Fig. 5D). To initially establish whether LRRK2 kinase inhibition influences Ad5 infectivity in the brain, we subjected *LRRK2* knockout (KO) and wild-type (WT) littermate adult mice to intrastriatal delivery of Ad5-eGFP vector at 4 distinct sites (∼4.2 x 10^9^ vp/site). At 10 days post-injection we observe equivalent levels of GFP in the striatum and SNpc of WT and KO mice, confirming that endogenous LRRK2 is not required for efficient Ad5 infection in the rodent brain (Fig. S7).

**Figure 5.**
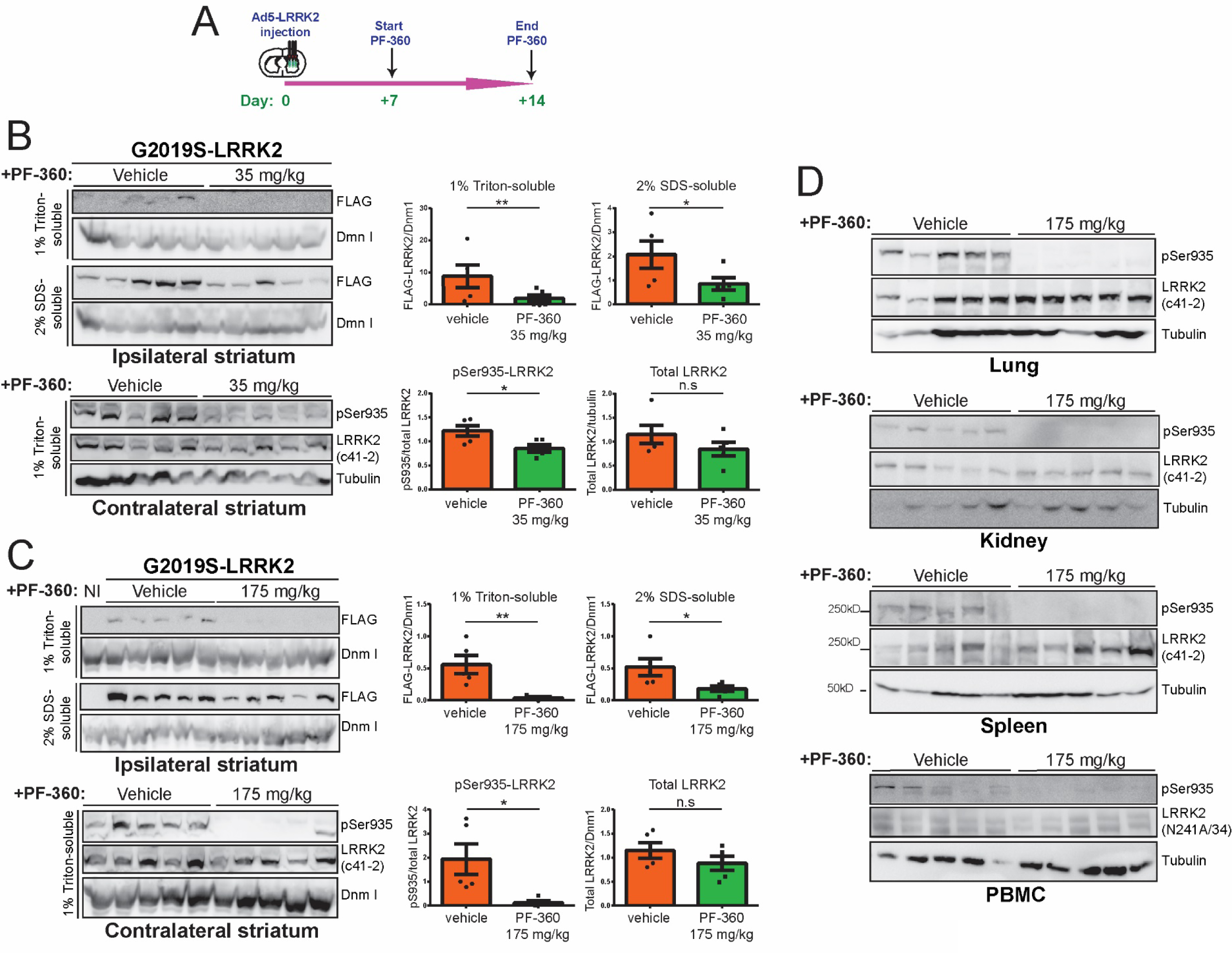
In-diet pharmacological kinase inhibition (PF-360) markedly destabilizes human G2019S-LRRK2 protein in rat brain. **A)** Ad5-G2019S-LRRK2 vectors were unilaterally delivered at six distinct locations (1.5 x 10^10^ vp/site, in 2.5 μl) in the rat striatum. Rats were fed with control (vehicle) or PF-360 (35 or 175 mg/kg) chow at day 7 post-injection for 7 consecutive days. Brain, peripheral tissues (lung, kidney, spleen) and PBMCs were collected at 14 days post-injection and subjected to sequential extraction in buffers containing 1% Triton X100 (Triton-soluble fraction) and 2% SDS (SDS-soluble fraction). **B-C)** Western blot analysis of ipsilateral striatum extracts indicating human G2019S-LRRK2 levels (anti-FLAG antibody), or contralateral striatum extracts indicating phosphorylated (pSer935) and total (MJFF2/c41-2) endogenous LRRK2 levels in rats fed with vehicle or PF-360 chow at **B)** 35 or **C)** 175 mg/kg chow. Dynamin I (DnmI) or β-tubulin were used as loading controls for normalization. NI, non-injected striatum. Densitometric quantitation is shown for human G2019S-LRRK2 (FLAG) levels in each detergent fraction, and endogenous pSer935-LRRK2 or total LRRK2 levels. Bars represent the mean ± SEM (*n* = 5 animals/vector), \**P*<0.05 or \*\**P*<0.01 by unpaired Student *t*-test. n.s. non-significant. **D)** Western blot analysis of peripheral tissues (lung, kidney and spleen) or PBMCs indicating levels of phosphorylated (pSer935) and total (c41-2 or N241A/34) endogenous LRRK2 in rats fed with vehicle or PF-360 (175 mg/kg) chow.

We next subjected rats to similar intrastriatal injections of Ad5-LRRK2 G2019S vector followed by chronic in-diet dosing of PF-360 (175 mg/kg) from 7 to 42 days post-injection to evaluate its neuroprotective effects (Fig. 6A). PF-360 dosing for 35 days results in a near-complete inhibition of endogenous LRRK2 kinase activity (pSer935) and the robust destabilization of human G2019S LRRK2 protein in the brain (Fig. S8A-C). PF-360 dosing is well-tolerated over this period with consistent and equivalent weekly body weight gain and food intake among treated animals (Fig. S8D-E), with rats receiving an estimated dose of ∼16 mg/kg/day PF-360. At 42 days post-injection, G2019S LRRK2 is robustly detected in the striatum and SNpc of animals treated with vehicle chow whereas PF-360 dosing leads to a marked reduction of FLAG-LRRK2 immunoreactivity (Fig. 6B). Hexon capsid immunostaining indicates equivalent levels of Ad5-LRRK2 particles in the striatum with vehicle or PF-360 treatment (Fig. 6D), further indicating that LRRK2 kinase inhibition does not compromise Ad5 infectivity. Consistent with this observation, PF-360 treatment does not alter the levels of reactive gliosis (Iba1 and GFAP) in the striatum induced by Ad5-LRRK2 delivery (Fig. 6D). Unbiased stereology reveals robust nigral dopaminergic neurodegeneration induced by G2019S LRRK2 (47.2 ± 4.3% neuronal loss) that is significantly attenuated by PF-360 administration (26.1 ± 3.1% loss) (Fig. 6B-C). Collectively, these data demonstrate that in-diet dosing with the LRRK2 kinase inhibitor, PF-360, provides neuroprotection against G2019S LRRK2 via an unexpected mechanism involving the selective destabilization of human LRRK2 protein.

**Figure 6.**
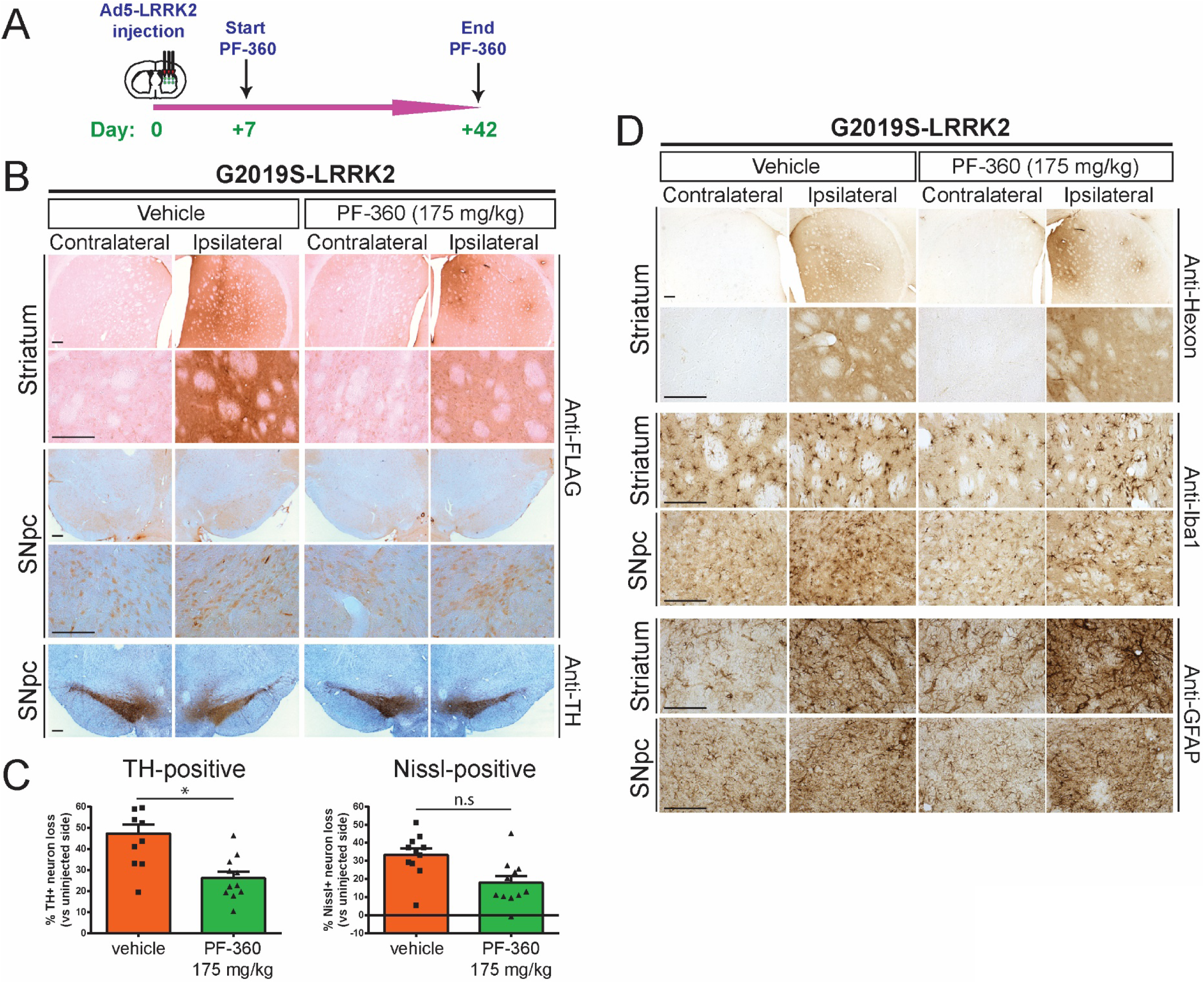
Pharmacological kinase inhibition (PF-360) protects against dopaminergic neurodegeneration induced by human G2019S LRRK2. **A)** Ad5-G2019S-LRRK2 vectors were unilaterally delivered at six distinct locations (1.5 x 10^10^ vp/site, in 2.5 μl) to the rat striatum. Rats were fed continuously with control (vehicle) or PF-360 (175 mg/kg) chow from days 7 to 42 post-injection. Brain tissues were harvested at 42 days post-injection and subjected to immunohistochemical analysis. **B)** Immunohistochemistry reveals expression of human LRRK2 (anti-FLAG antibody) in striatum and nigra, or nigral dopaminergic neurons (anti-TH antibody), of rats at 42 days post-injection. Images indicate low and high magnification for each group and brain region. TH sections were counterstained with Cresyl violet. Scale bars: 500 μm. **C)** Unbiased stereological analysis of TH-positive dopaminergic and total Nissl-positive neuron number in the substantia nigra at 42 days post-injection. Bars represent % neuronal loss in the injected ipsilateral nigra relative to the contralateral nigra (mean ± SEM, *n* = 12 animals/group), \**P*<0.05 by unpaired Student *t*-test. n.s. non-significant. **D)** Immunohistochemistry at 42 days post-injection showing expression of Ad5 capsid (anti-Hexon) in the ipsilateral striatum, or microglia (Iba1) and astrocytes (GFAP) in striatum and substantia nigra of rats expressing human G2019S-LRRK2 treated with vehicle or PF-360 chow. Scale bars: 500 μm.

### Contribution of GTPase activity to G2019S LRRK2-induced neurodegeneration in rats

GTP-binding via the phosphate-binding loop (P-loop) can increase the kinase activity of LRRK2 (13, 14, 16), whereas synthetic mutations that disrupt P-loop GTP-binding (K1347A or T1348N) markedly impair GTP hydrolysis and kinase activity (13, 14). PD-linked mutations in the Roc-COR tandem domain commonly impair the GTP hydrolysis activity of LRRK2 thereby prolonging the GTP-bound ‘on’ state and indirectly enhancing kinase activity (15). We have previously identified hypothesis-testing mutations in the Switch II catalytic motif of the Roc domain that markedly enhance (R1398L) or reduce (R1398L/T1343V) GTP hydrolysis activity *in vitro* and lead to an increase in the GDP-bound ‘off’ or GTP-bound ‘on’ state of LRRK2, respectively (13, 23). Given the importance of the GTPase domain to LRRK2 activity and the paucity of selective small molecules targeting this domain, we sought to employ these genetic mutations to evaluate the contribution of GTP-binding and GTP hydrolysis activity to neurodegeneration induced by G2019S LRRK2 in the rat brain. Accordingly, new high-titer Ad5-LRRK2 G2019S vectors were developed that additionally harbor distinct functional mutations in the Roc domain (G2019S/T1348N, G2019S/R1398L and G2019S/R1398L/T1343V). To initially evaluate these Ad5-LRRK2 vectors, we demonstrate the titer-dependent and equivalent expression levels of G2019S, G2019S/R1398L and G2019S/R1398L/T1343V LRRK2 variants in SH-SY5Y cells by Western blot analysis (Fig. 7A) whereas G2019S/T1348N LRRK2 levels are reduced indicating the impaired stability of this variant. The levels of phosphorylation (Ser935, Ser910 and Ser1292) of G2019S/R1398L and G2019S/R1398L/T1343V LRRK2 are similar to those of G2019S LRRK2 suggesting that modulating GTPase activity does not alter the elevated kinase activity of G2019S LRRK2 in cells (Fig. 7A), similar to our prior *in vitro* studies (13). G2019S/T1348N LRRK2 exhibits a marked reduction in phosphorylation at these three sites indicating impaired kinase activity (Fig. 7A), similar to *in vitro* data (13), confirming that GTP-binding capacity is critical for normal kinase activity (13, 14, 16). These data demonstrate that genetic inhibition of GTP-binding impairs the protein stability and kinase activity of G2019S LRRK2.

**Figure 7.**
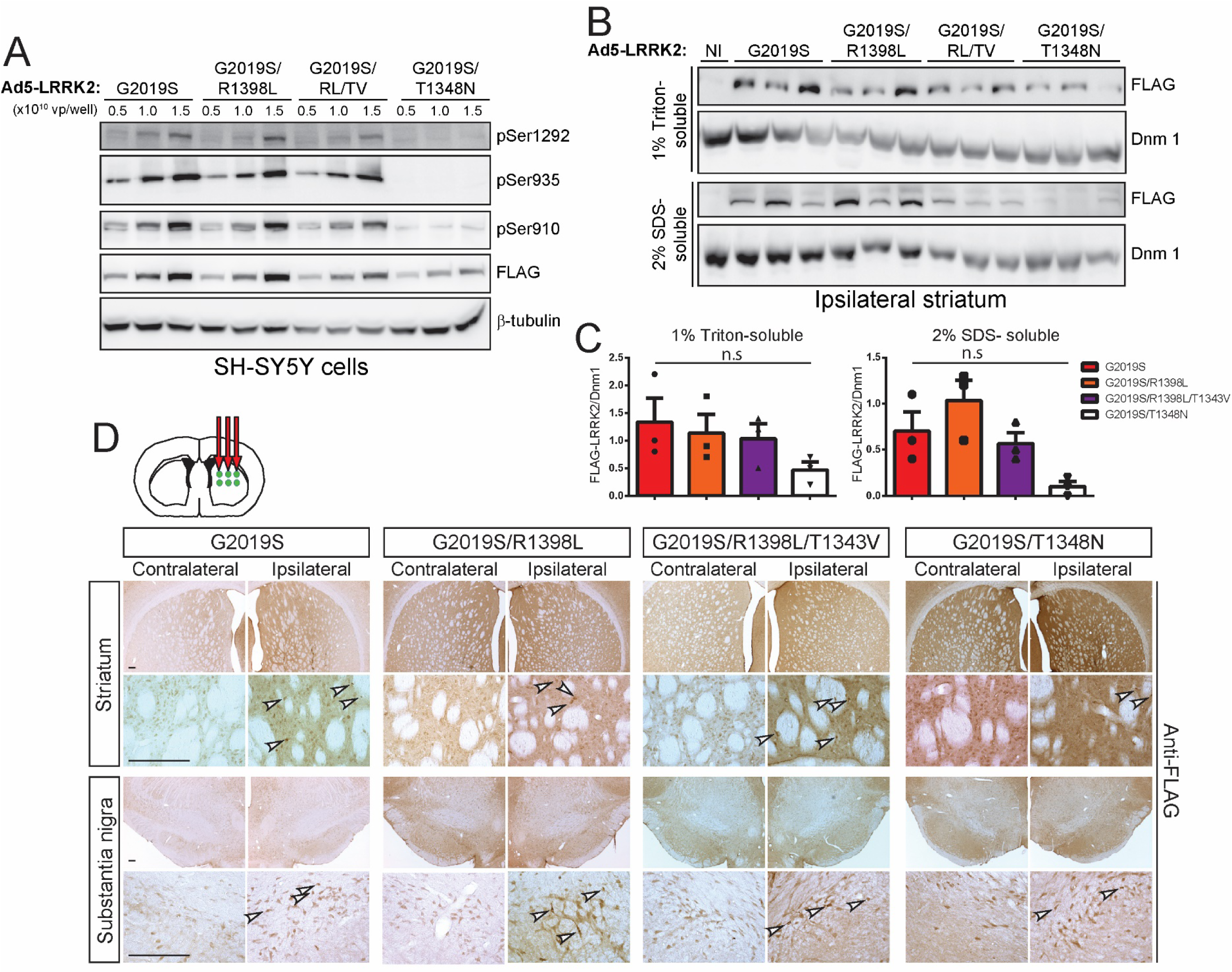
Evaluation of Ad5-LRRK2-G2019S vectors with genetic disruption of GTP-binding or GTP hydrolysis in cells and rat brain. **A)** Validation of high-titer Ad5-LRRK2 (G2019S, G2019S/R1398L, G2019S/R1398L/T1343V or G2019S/T1348N) vectors and LRRK2 phosphorylation in SH-SY5Y cells. Cells were transduced with increasing viral particles (vp) of each Ad5-LRRK2 vector (0.5, 1 or 5.0 x 10^10^ vp/well). Western blot analysis of cell extracts with FLAG antibody (transduced human LRRK2) or antibodies to detect LRRK2 autophosphorylation (pSer1292) or constitutive phosphorylation (pSer935, pSer910) sites to detect a titer-dependent increase in human LRRK2 levels/activity. β-tubulin was used as protein loading control. **B)** Ipsilateral striatum was harvested at 10 days after unilateral intrastriatal delivery of Ad5-LRRK2 vectors and subjected to sequential detergent extraction in 1% Triton X100 and 2% SDS. Striatal fractions were analyzed by Western blot analysis (*n* = 3 animals/vector) indicating human LRRK2 variant levels (anti-FLAG antibody). Expression of dynamin-I (DnmI) was used as loading control. NI, non-injected striatum. **C)** Densitometric analysis of human LRRK2 variant levels (FLAG) normalized to Dnm 1 levels in each detergent fraction. Bars represent the mean ± SEM (*n* = 3 animals/vector). *n.s*, non-significant by one-way ANOVA with Bonferroni’s multiple comparisons. **D)** Ad5-LRRK2 (G2019S, G2019S/R1398L, G2019S/R1398L/T1343V, G2019S/T1348N) vectors were unilaterally delivered at six injection sites (1.5 x 10^10^ vp/site, in 2.5 μl) in the rat striatum. Immunohistochemistry indicates human LRRK2 variant (anti-FLAG) expression in the ipsilateral striatum and substantia nigra that persists up to 42 days post-injection. Images indicate low and high magnification for each vector and brain region. Scale bars: 500 μm.

To explore the effects of these GTPase mutations *in vivo*, we delivered Ad5-LRRK2 vectors (G2019S, G2019S/R1398L, G2019S/R1398L/T1343V and G2019S/T1348N) to the striatum of rats (∼1.25 x 10^10^ vp/site) and conducted biochemical analysis at 10 days post-injection. Human LRRK2 variants are detected in 1% Triton-soluble and -insoluble extracts from the ipsilateral striatum with equivalent levels of G2019S, G2019S/R1398L and G2019S/R1398L/T1343V LRRK2 proteins by Western blot and densitometric analyses (Fig. 7B-C). The levels of G2019S/T1348N LRRK2 are dramatically reduced in the striatum relative to other LRRK2 variants, further supporting the reduced protein stability of this GTPase variant (Fig. 7B-C). Our data indicate that functional mutations that alter GTP hydrolysis are well-tolerated in cells and *in vivo* but do not alter the kinase activity of G2019S LRRK2, whereas inhibition of GTP-binding compromises LRRK2 protein stability and kinase activity, consistent with prior studies (13, 14, 16).

The neuroprotective capacity of these functional GTPase variants was next evaluated at 42 days following the intrastriatal delivery of Ad5-LRRK2 vectors. FLAG-LRRK2 immunoreactivity indicates that G2019S, G2019S/R1398L and G2019S/R1398L/T1343V LRRK2 proteins are detected in the ipsilateral striatum and SNpc at equivalent levels whereas G2019S/T1348N LRRK2 levels are markedly reduced (Fig. 7D). Hexon capsid immunostaining indicates equivalent levels of Ad5 particles for each LRRK2 variant in the striatum (Fig. 8A), indicating that G2019S/T1348N LRRK2 exhibits reduced protein stability in the brain. Intriguingly, unbiased stereology reveals robust dopaminergic neuronal loss induced by G2019S LRRK2 (33.4 ± 8.8% loss) that is significantly attenuated with G2019S/R1398L (17.9 ± 8.8%) or G2019S/T1348N (9.8 ± 7.3%) LRRK2 variants (Fig. 8B-C). Neuronal loss induced by G2019S/R1398L/T1343V LRRK2 (24.6 ± 11.9% loss) is also reduced relative to G2019S LRRK2 although this effect is not significant (Fig. 8B-C). The loss of total Nissl-positive SNpc neurons parallels the loss of TH-positive SNpc neurons in this rat model thereby confirming neuronal cell death (Fig. 8C). Taken together, these data suggest that genetically enhancing GTP hydrolysis (R1398L) or disrupting GTP-binding (T1348N) are sufficient to protect against G2019S LRRK2 in the rat brain. However, the protective effects elicited by the T1348N variant are likely mediated by destabilizing the LRRK2 protein rather than via diminished GTP-binding *per se*.

**Figure 8.**
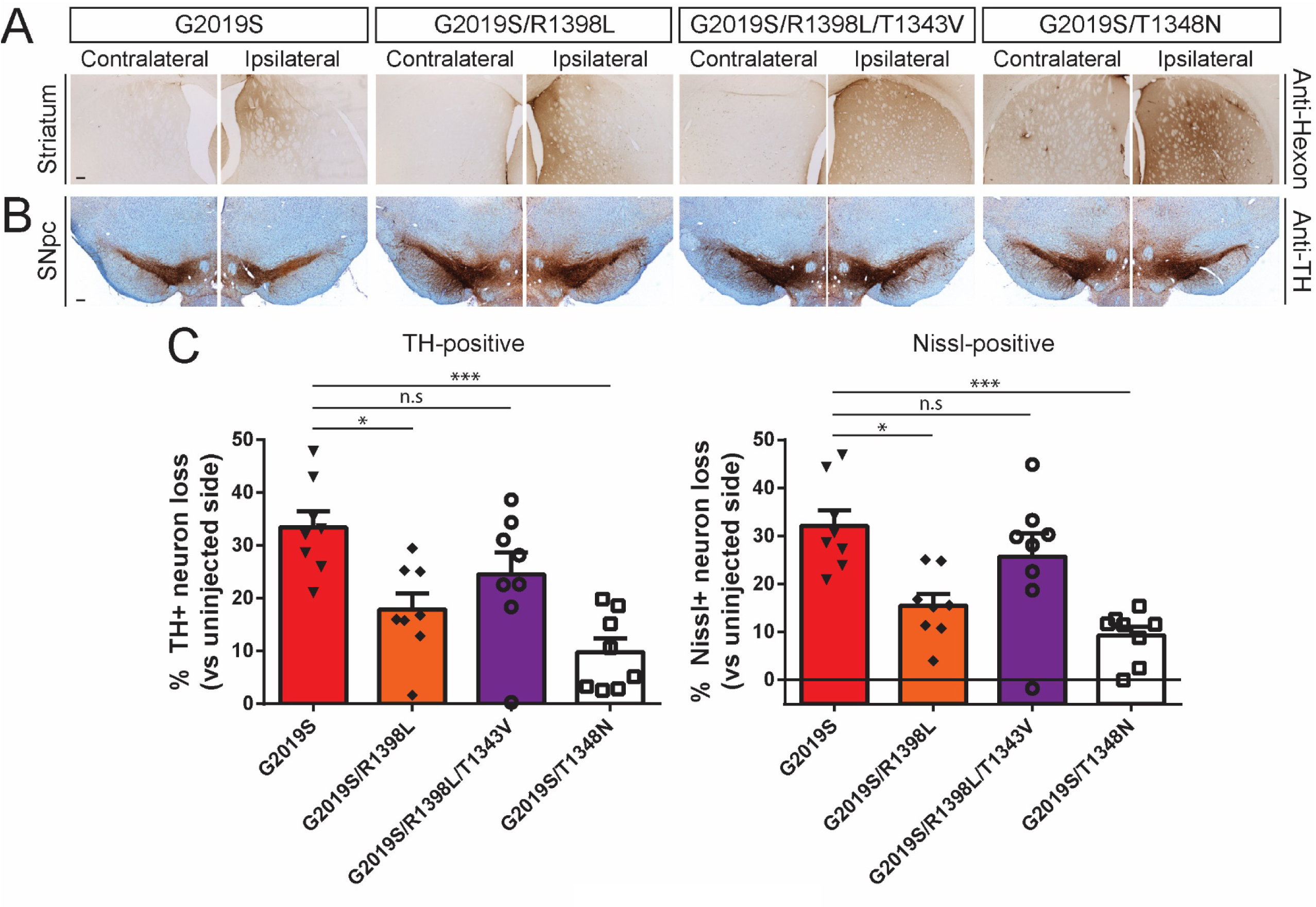
Ad5-G2019S-LRRK2 induces dopaminergic neurodegeneration in a GTPase-dependent manner in adult rat brain. **A)** Immunohistochemistry indicating Ad5 capsid (anti-Hexon) in the ipsilateral rat striatum at 42 days post-injection of Ad5-LRRK2 vectors. Scale bars: 500 μm. **B)** Immunohistochemistry showing nigral dopaminergic neurons (anti-TH antibody) at 42 days post-delivery. All sections were counterstained with Cresyl violet. Scale bars: 500 μm. **C)** Unbiased stereological analysis of TH-positive dopaminergic and total Nissl-positive neuron number in the substantia nigra at 42 days post-injection. Bars represent % neuronal loss in the injected ipsilateral nigra relative to the contralateral nigra (mean ± SEM, *n* = 8 animals/vector), \**P*<0.05 or \*\*\**P*<0.001 by one-way ANOVA with Bonferroni’s multiple comparisons test. *n.s*, non-significant.

## Discussion

Transgenic and knockin rodent models of *LRRK2*-linked PD generally exhibit mild phenotypes which has hampered the identification of key mechanisms that drive neurodegeneration due to familial mutations (12, 33). However, prior studies using large-capacity viral vectors, such as HSV or Ad5, delivered to the brains of adult rodents have proven successful in recapitulating the age-dependent degeneration of substantia nigra dopaminergic neurons induced by the common G2019S mutation in LRRK2 (29, 37). Our prior studies using Ad5-LRRK2 vectors suggested that G2019S LRRK2 induces neuropathology in rats via a kinase-dependent mechanism (32). However, these pathological effects were confined to the striatum and relied upon a kinase-inactive version of the LRRK2 protein, D1994N, that is unstable in rat brain and primary neurons (13, 32). In the present study, we further develop and optimize this Ad5-LRRK2 rat model of PD to demonstrate that human G2019S LRRK2 induces robust nigral dopaminergic neurodegeneration over 6 weeks following the stereotactic delivery of Ad5 vectors directly to the striatum or substantia nigra. Using a stable kinase-inactive mutation in the ATP-binding pocket, K1906M, we show that G2019S LRRK2 induces neuronal loss through a kinase-dependent mechanism. In addition, we demonstrate that the pharmacological inhibition of LRRK2 kinase activity using the selective and potent inhibitor PF-360 can attenuate dopaminergic neuronal loss induced by G2019S LRRK2 in rat brain. Unexpectedly, PF-360-mediated kinase inhibition selectively destabilizes the exogenous human LRRK2 protein in the rat brain relative to its modest effects on endogenous LRRK2. Our data also reveal for the first time a critical role for GTPase activity in mediating the pathogenic actions of G2019S LRRK2. Intriguingly, a mutation in the Switch II motif of the Roc domain that enhances GTP hydrolysis activity, R1398L, attenuates neuronal loss induced by G2019S LRRK2 via a mechanism that may not involve kinase activity. A second mutation that impairs GDP/GTP-binding via the Roc P-loop motif, T1348N, is also neuroprotective albeit by dramatically destabilizing the LRRK2 protein. Taken together, these data convincingly support a critical role for kinase and GTPase activity in mediating the neurodegenerative effects of the LRRK2 G2019S mutation *in vivo*.

It has proven difficult to develop relevant and robust preclinical animal models for PD-linked *LRRK2* mutations to directly explore the underlying mechanisms of neurodegeneration and evaluate novel therapeutic strategies (12, 33). Viral-mediated gene transfer for human LRRK2 has allowed the identification of robust neuropathology in rodent models although subsequent mechanistic studies have been lacking. Although HSV vectors were initially used to reveal dopaminergic neuronal loss induced by human G2019S LRRK2 in the mouse brain, a role for kinase activity in this model relied upon an unstable kinase-inactive mutation (D1994A) and the use of generation “0” LRRK2 inhibitors that are not selective for LRRK2 (i.e. GW5074 or indirubin-3’-monoxime) (29). For example, GW5074 is a potent inhibitor of c-Raf1 kinase with an IC_50_ of ∼9 nM, whereas indirubin-3’-monoxime potently inhibits GSK-3β (IC_50_ ∼22 nM), making it difficult to assign neuroprotective effects specifically to LRRK2 kinase inhibition in this model. Furthermore, the HSV-LRRK2 model no longer appears to be in active use. Our prior studies with neuronal-expressing Ad5-LRRK2 vectors in rat brain also revealed specific neuronal loss and neuropathology with G2019S LRRK2 but also employed an unstable kinase-inactive D1994N mutation in LRRK2 (32). A similar study using broadly-expressing Ad5-LRRK2 vectors in mouse brain compared G2019S and another kinase-inactive mutation, D1994A LRRK2, revealing neuroinflammation and vacuolization in aged mice due to LRRK2 expression in glial cells (48). A recent study in the non-human primate, *Microcebus murinus*, using long-acting canine adenovirus type 2 vectors, revealed an equivalent level of neuronal loss induced by WT and G2019S LRRK2 but without evaluating kinase activity (49). Therefore, the contribution of kinase activity in these or any mutant LRRK2 animal model has not been clearly established. So far, pharmacological LRRK2 kinase inhibition has been evaluated indirectly in rodent models with AAV-mediated human α-synuclein expression where PF-475 protects against dopaminergic neurodegeneration in rats induced by WT α-synuclein (50) but PF-360 lacked neuroprotection against A53T α-synuclein (51). Despite the paucity of direct evidence in LRRK2 animal models, selective LRRK2 kinase inhibitors (i.e. DNL151, DNL201) are currently in phase I clinical trials.

Our data extend these prior observations in LRRK2 viral models by firstly demonstrating robust dopaminergic neurodegeneration induced by G2019S LRRK2 in the rat brain when transgene expression is restricted to neurons using the human synapsin-1 promoter in the Ad5 vector. This finding implies that G2019S LRRK2 is able to induce dopaminergic neuronal loss via a cell-autonomous mechanism but does not exclude a non-cell-autonomous role for LRRK2 in astrocytes and/or microglia. Indeed, Ad5 vector delivery in this rat model induces modest and prolonged neuroinflammation which may itself contribute to or be required for the neurotoxic effects of G2019S LRRK2. Secondly, we demonstrate a key role for kinase activity in the Ad5-LRRK2 model using a kinase-inactive mutant (K1906M) that is stably expressed in rat brain relative to kinase-active LRRK2 variants. Importantly, by monitoring autophosphorylation at Ser1292 in cells and brain, we show that the K1906M mutation impairs the kinase activity of G2019S LRRK2. Our attempts to use pT73-Rab10 to monitor G2019S LRRK2 activity in rat brain were unsuccessful most likely because endogenous Rab10 and exogenous human LRRK2 reside in different cell populations in this model, and potentially because the G2019S mutation produces a more modest increase in Rab10 phosphorylation compared to other PD-linked familial mutations (18). Introducing the K1906M mutation robustly rescues nigral neuronal loss induced by G2019S LRRK2 following either intrastriatal or intranigral Ad5 delivery, although intrastriatal delivery produced lower background effects of the Ad5 vectors. Third, the genetic analyses of LRRK2 kinase activity were complemented by pharmacological studies with PF-360, a potent and selective generation “3” LRRK2 inhibitor. PF-360 is highly potent with an *in vivo* IC_50_ of ∼3 nM for LRRK2, and exhibits an improved selectivity profile, oral bioavailability and brain permeability compared to earlier generations of LRRK2 inhibitors (41, 47). We demonstrate full inhibition of endogenous LRRK2 kinase activity in peripheral tissues and brain of our rat model with chronic in-diet dosing of PF-360 at 175 mg/kg chow. While PF-360 produces neuroprotection against G2019S LRRK2 in rats at this dose, it most likely achieves this effect via an unexpected mechanism involving the selective and robust destabilization of human LRRK2 protein in brain compared to its mild effects on endogenous rat LRRK2. This destabilizing effect was also prominently observed at lower doses of PF-360 (35 mg/kg chow). Structurally distinct LRRK2 kinase inhibitors (i.e. MLi-2, PF-475, GSK2578215A, HG 10-102-01) have been shown to induce LRRK2 protein destabilization via proteasomal degradation in cells and tissues that is dependent on dose and exposure time (52). This effect is consistent with the destabilizing effects of certain kinase-inactive mutations (D1994A/N/S) (13, 26, 31, 32), suggesting an intricate role for kinase activity in maintaining normal stability and limiting the turnover of LRRK2 protein. However, our data imply a strong bias for destabilization either due to species (human over rat LRRK2), mutational status (G2019S over wild-type LRRK2) or perhaps due to levels of overexpression and availability (exogenous over endogenous LRRK2). These findings could have ramifications for the treatment of humans with LRRK2 kinase inhibitors, although it is not feasible at present to determine whether kinase inhibition will dramatically destabilize brain LRRK2 or show a preference for mutant over wild-type LRRK2 in human tissues. In any case, the destabilizing effect of PF-360 treatment on G2019S LRRK2 in our model proves target engagement and indicates that early inhibition and/or reduction of mutant LRRK2 in this model is sufficient for neuroprotection. Our future studies will explore the impact of distinct kinase inhibitors in the Ad5-LRRK2 rat model and whether LRRK2 destabilization is mutation-specific.

The contribution of GTPase activity in LRRK2 animal models has not been studied, largely owing to a lack of robust activity assays and pharmacological tools. While LRRK2 has been shown to bind GTP and exhibit a slow rate of GTP hydrolysis *in vitro* (13, 14, 16, 20–23), comparable activity assays in tissues are lacking. Putative pharmacological inhibitors of LRRK2 GTP-binding have also been reported (i.e. FX2149) although their selectivity profiles and mode of action are poorly defined (53, 54). At present, mutational analysis of the Roc GTPase domain remains the only reliable method to probe its function *in vivo*. We have previously identified hypothesis-testing mutations in conserved motifs that regulate the GTPase activity of LRRK2 (13, 23). In our hands, the Switch II motif mutation, R1398L, increases GTP hydrolysis *in vitro* but has no obvious effect on steady-state GTP-binding (13, 23). In combination with a second P-loop mutation, R1398L/T1343V, this variant impairs GTP hydrolysis *in vitro* (13). Disruption of the P-loop motif via T1348N impairs GDP/GTP-binding and therefore prevents GTP hydrolysis (13). Based on these *in vitro* studies, we are able to demonstrate that the R1398L and T1348N mutations provide robust neuroprotection against G2019S LRRK2 in the rat brain but perhaps via different mechanisms. While the G2019S/R1398L protein is stable *in vivo*, the G2019S/T1348N protein is highly unstable and therefore phenocopies a loss-of-function effect similar to the effects of kinase inhibitors (13). The data suggest to us instead that GTP-binding capacity is critical for maintaining the normal stability of the LRRK2 protein *in vivo* although the underlying mechanism is unclear. We have previously shown that T1348N LRRK2 is unstable, interacts poorly with itself, and is enzymatically inactive *in vitro* and in cells (13), potentially consistent with impaired dimerization. For the R1398L mutation that is normally dimeric in cells (13), we assume its protective effect *in vivo* is due to increased GTP hydrolysis activity but it could also disrupt the GTPase cycle of LRRK2 in general. Unlike the effects of K1906M, the R1398L mutation does not alter the elevated kinase activity of G2019S LRRK2 in cells, suggesting that neuroprotection *in vivo* is most likely mediated via a kinase-independent mechanism. Nevertheless, our data support interference with GTP hydrolysis via the Switch II motif or GTP-binding via the P-loop as promising neuroprotective strategies for alleviating the pathogenic effects of G2019S LRRK2. The underlying mechanism(s) involved and whether these protective effects extend to PD-linked *LRRK2* mutations outside of the kinase domain (i.e. N1437H, R1441C/G/H, Y1699C) await further study.

Collectively, our data provide a comprehensive evaluation of Ad5-LRRK2 vectors for exploring the neurodegenerative effects and mechanisms of PD-linked *LRRK2* mutations in neurons of the adult rat brain. While dopaminergic neuronal loss provides a robust phenotypic readout of G2019S LRRK2 in this model, we identify axonal damage as another specific phenotype, and also define non-specific neuropathological markers that are not informative for LRRK2 pathophysiology. As a proof-of-concept we demonstrate that this Ad5-LRRK2 model can be effectively used to evaluate the genetic and pharmacological contribution of enzymatic activity to LRRK2-dependent neurodegeneration. We nominate kinase inhibition, increased GTP hydrolysis and inhibition of GTP-binding as potential therapeutic strategies for attenuating the neurotoxic effects of LRRK2 mutations. The Ad5-LRRK2 rat model provides a relatively robust, flexible and relevant animal model for rapidly exploring novel mechanisms and therapeutic strategies in PD.

## Materials and Methods

### Animals

Adult female Wistar rats (180-200 g) were obtained from Harlan Laboratories. *LRRK2* KO mice with a deletion of exon 41 (31) were kindly provided by Drs. Giorgio Rovelli and Derya Shimshek (Novartis Pharma AG, Basel Switzerland). Rodents were maintained in a pathogen-free barrier facility and provided with food and water *ad libitum*, and exposed to a 12 h light/dark cycle. Animals were treated in strict accordance with the NIH Guide for the Care and Use of Laboratory Animals. All animal experiments were approved by the Van Andel Institute Institutional Animal Care and Use Committee (IACUC).

### Cell culture

Human SH-SY5Y neuroblastoma cells (ATCC) were maintained in Dulbecco’s modified Eagle’s media (DMEM) with high glucose and without pyruvate supplemented with 10% fetal bovine serum and 1X penicillin/streptomycin at 37°C in a humidified atmosphere containing 5% CO_2_. Stable E2T-293 cells used for adenovirus production were maintained in DMEM with 10% fetal bovine serum, 1X GlutaMAX (Thermo Fisher), G418 (200 μg/ml), puromycin (1 μg/ml; Sigma), tetracycline (1 μg/ml; Sigma) and 1X penicillin/streptomycin. Cells were routinely passaged at intervals of 3-4 days and culture was terminated after 25 passages.

### Adenovirus production

Recombinant E1, E2a, E3-deleted (second generation) human adenovirus serotype 5 (Ad5) were produced and purified as previously described (32, 37). Codon-optimized full-length WT and G2019S human LRRK2 cDNAs, tagged at the N-terminus with a 3xFLAG epitope were cloned into a modified pDC511 shuttle plasmid downstream of a human synapsin-1 promoter and synthetic intron (37, 55). A D1994N mutation was introduced into the pDC511-G2019S LRRK2 by site-directed mutagenesis using the QuickChange XL kit (Stratagene) as described (32). Similarly, K1906M, R1398L and T1348N mutations were introduced into pDC511-G2019S LRRK2, and the T1343V mutation into pDC511-G2019S/R1398L LRRK2 by site-directed mutagenesis. DNA sequencing across the entire LRRK2 cDNA sequence confirmed the presence of each mutation and the absence of any secondary mutations.

To generate Ad5 vectors, human Ad5 genomic plasmid (pBHGfrtΔE1,3FLP; Microbix) and each pDC511-LRRK2 shuttle plasmid were introduced by calcium phosphate (Invitrogen) transfection into stable Tet-On-inducible E2a cells (E2T cells) that express the complementing E2a (adenovirus early gene) in the presence of tetracycline (56). To maintain consistency between each virus production, E2T cells at an early passage number (P3-P6) were used for transfection. The adenoviral vector results from FLP-mediated site-specific recombination between these two plasmids. Cells were harvested at 14 days post-transfection upon becoming cytopathic and were subjected to lysis by three consecutive freeze-thaw cycles. Serial dilutions (10^0^ to 10^-5^) of viral supernatants were used to infect E2T cells (5 x 10^5^ cells/well in 6-well plate) for plaque assays and cells were maintained in culture medium containing 5% SeaPlaque agarose (Lonza) until plaques become visible at ∼12 days after infection. In general, we collected 5 isolated plaques per virus that were subjected to sequential amplification steps in 24-well, 6-well and 10 cm dishes, respectively. When cells become cytopathic after amplification in 10 cm dishes, they were harvested and subjected to lysis. Supernatants from each viral clone were used to infect SH-SY5Y cells for expression screening by Western blot analysis. Viral clones inducing equivalent LRRK2 expression were selected for final amplification in 15 cm dishes. After infection, cells became cytopathic by ∼7 days and were harvested and lysed. Total cell supernatants were filtered over PES membranes (0.45 μm; Millipore) and stored as “infective medium” at −80°C for further use.

To prepare high titer viral stocks, infective medium for each virus was used to infect E2T cells plated in 15 cm dishes (20 culture dishes/virus). For consistency, only E2T cells at early passage number (P3-P6) were used for virus amplification. At 7 days post-infection when cells become cytopathic, they were harvested and lysed. Total supernatants were treated with benzonase nuclease (12.5 U/ml) and subjected to two rounds of column purification using the Vivapure AdenoPACK 100RT kit (Sartorius). The final viral stocks were stored at −80°C in storage solution (10 mM HEPES, 250 mM sucrose, 1 mM MgCl_2_ pH 7.4). The concentration of viral particles was determined by spectrophotometry using OD_260_ measurements.

### Adenovirus infectivity: SH-SY5Y cells

SH-SY5Y cells were infected with each purified adenovirus to verify equivalent LRRK2 expression. In general, 5 x 10^5^ cells/ virus dose were plated in 35 mm dishes 24 h before infection. On the day of infection, purified viruses were mixed at the desired volumes in 0.5 ml of normal culture media. Cells were incubated with virus at 37°C with gentle rocking every 10 min. After 30 min of incubation, 1.5 ml of fresh media was added to each dish and cells were incubated at 37°C. At 48 h post-infection, cells were washed twice in cold TBS (50 mM Tris pH 7.5, 150 mM NaCl) and harvested in 200 μl lysis buffer (50 mM Tris pH 7.5, 150 mM NaCl, 1% Triton-X100, 1X Complete protease inhibitor cocktail [Roche], 1X phosphatase inhibitor cocktails 2 and 3 [Sigma]). Protein concentration was determined using BCA assays (Pierce Biotech).

### Stereotactic rodent surgery

Adult female Wistar rats (Harlan Laboratories) weighing approximately 180-200 g, or adult homozygous *LRRK2* KO and wild-type littermate mice (mixed sex), were subjected to stereotactic surgery for unilateral delivery of recombinant adenoviral vectors. After the animals were deeply anesthetized with continuous isofluorane, they were placed in a stereotactic frame (Model 963-A, David Kopf Instruments) and the cranium was exposed. For intranigral injection, viral vectors were delivered at a single point in the right SNpc using the following coordinates relative to bregma: anterior-posterior −5.2 mm, mediolateral −2 mm, and dorsoventral −7.8 mm relative to the skull surface. Blunt tip steel needles (34-gauge) connected through polyethylene tubing to a 10 μl Hamilton syringe were inserted slowly to the injection site and left in place for 30 s before starting the infusion. A volume of ∼2.5 μl of virus was injected at a flow rate of 0.2 μl/min with an automatic pump (CMA Microdialysis). The needle was left in place for an additional 5 min before being slowly withdrawn to allow the tissue to close over and minimize leakage of the injected virus. The skin was sealed by stapling and the animals were monitored until they recovered from anesthesia. Staples were removed at 10 days post-injection. The animals were sacrificed at 21 days post-surgery.

For intrastriatal injections, viral vectors were delivered at 6 points in the right dorsal striatum (2 dorsoventral deposits along three needle tracks) of rats using the following coordinates relative to bregma: anterior-posterior +0.48 mm, mediolateral −2 mm, −3 mm and −4 mm, and dorsoventral −6 mm and −4.8 mm relative to the skull surface. Volumes of ∼2.5 μl of virus was injected per site, and needles were left in place for 5 min between the first and second infusions (at DV, −6 and −4.8 mm), and for an additional 5 min after the second infusion before being withdrawn. Rats were sacrificed at 10 or 42 days post-surgery. For mice, viral vectors (1 μl/site) were delivered at 4 points in the right dorsal striatum (2 dorsoventral deposits along two needle tracks) using the coordinates: anterior-posterior +0.86 mm, mediolateral −1.2 mm and −2.2 mm, and dorsoventral −3.7 mm and −3.0 mm relative to the skull surface. Mice were sacrificed at 10 days post-surgery.

### Immunohistochemistry

Animals were perfused transcardially with saline followed by 4% paraformaldehyde (PFA) in 0.1 M phosphate buffer (pH 7.3). After post-fixation for 24 h in 4% PFA and cryoprotection in 30% sucrose solution, brains were dissected into 40 μm-thick coronal sections using a sliding microtome (SM2010R, Leica). For chromogenic immunostaining, sections were washed 3x in TBS then blocked for endogenous peroxidase activity by incubation in 3% H_2_O_2_ (Sigma) diluted in methanol for 10 min at 4°C. Sections were blocked for non-specific binding in TBS containing 10% normal goat serum (Invitrogen) supplemented with 0.1% Triton-X100 for 1 h at room temperature. Sections were incubated in primary antibodies for 48 h at 4°C and biotinylated secondary antibodies (Vector Labs) for 2 h at room temperature. After incubation with ABC reagent (Vector Labs) for 1 h at room temperature and visualization in 3,3’-diaminobenzidine tetrahydrochloride (DAB; Vector Labs), sections were mounted on Superfrost plus slides (Fisher Scientific), dehydrated with increasing ethanol concentrations and xylene, and coverslipped using Entellan (Merck). Images were captured using a light microscope (Axio Imager M2, Zeiss) equipped with a color CCD camera (AxioCam, Zeiss).

For immunofluorescence staining, sections were washed 3x in TBS then blocked for non-specific binding in TBS containing 10% normal goat serum (Invitrogen) supplemented with 0.1% Triton-X100 for 1 h at room temperature. Sections were incubated in primary antibodies for 48 h at 4°C and secondary antibodies conjugated to AlexaFluor-488 or AlexaFluor-594 (Life Technologies) for 2 h at room temperature. Sections were washed in TBS with 0.1% Tween 20, mounted on Superfrost plus slides (Fisher Scientific) and coverslipped using Prolong mounting medium containing DAPI (Invitrogen). Sections were imaged using a Nikon A1plus-RSi scanning confocal microscope.

For Gallyas silver staining, sections were pre-incubated in 4% PFA (in 0.1 M PB) for ≥2 days at 4°C before processing with the FD NeuroSilver™ kit II (FD Neurotechnologies, Inc.) according to the manufacturer’s protocol. Sections were mounted on Superfrost plus slides (Fisher Scientific), dehydrated with increasing ethanol concentrations and xylene, and coverslipped using Entellan (Merck). Images were captured using a light microscope (Axio Imager M2, Zeiss) equipped with a color CCD camera (AxioCam, Zeiss).

### Brain tissue fractionation

Brain tissues for biochemical analyses were harvested at 10 or 42 days after Ad5 delivery. Striatum and ventral midbrain from injected and non-injected hemispheres were rapidly dissected and homogenized with 6 volumes of Triton lysis buffer (50 mM Tris pH 7.5, 150 mM NaCl, 5% glycerol, 1% Triton-X100, 1 mM EDTA, 1X EDTA-free Complete protease inhibitor cocktail [Roche] and 1X phosphatase inhibitor cocktail 2 and 3 [Sigma]). Tissues were disrupted using a mechanical homogenizer (IKA® T10 basic, Ultra Turrax). The Triton-soluble fraction was obtained after ultracentrifugation at 100,000*g* for 30 min at 4°C. The resulting pellets were further extracted by sonication in 3X volumes of SDS lysis buffer (50 mM Tris-HCl pH 7.4, 2% SDS, 1X EDTA-free Complete protease inhibitor cocktail [Roche] and 1X phosphatase inhibitor cocktail 2 and 3 [Sigma]). Samples were centrifuged at 21,000*g* for 30 min at 25°C to obtain the Triton-insoluble (SDS-soluble) fraction. Protein concentration was determined using the BCA assay (Pierce Biotech).

### In-diet PF-360 dosing and pharmacodynamic assays

PF-360 was synthesized as described in patent US2014/005183 at >98% purity and formulated into medicated chow at 35 or 175 mg/kg by Research Diets, as previously described (41). PF-360 chow was kindly provided by Dr. Andrew West (Duke University). Animals were fed chow continuously for 7 or 35 days beginning at 7 days after stereotactic surgery. Body weight and chow consumption was monitored every 7 days. Pharmacodynamic assays to monitor LRRK2 kinase inhibition by in-diet PF-360 dosing included assessment of endogenous pSer935 levels normalized to total LRRK2 levels in brain and peripheral tissues (lung, kidney, spleen, PBMCs). PBMCs were isolated from tail vein blood (500 µl) using SepMate-15 tubes (STEMCELL Technologies).

### Western blot analysis

Equivalent protein samples were mixed with 2X Laemmli sample buffer (Bio-Rad) containing 5% 2-mercaptoethanol and resolved by SDS-PAGE using 7.5% Tris-Glycine or 4-20% Tris-Glycine gels (Novex) followed by electrophoretic transfer onto nitrocellulose membranes (0.2 μm, Amersham). Membranes were blocked in TBS with 5% non-fat milk (Bio-Rad) for 1 h at room temperature, and incubated with primary antibodies overnight at 4°C. For probing with multiple antibodies, membranes were sequentially stripped in Restore™ Western Blot buffer (Thermo Scientific) for 30 min at 37°C and re-blocked for 1 h at room temperature. Secondary antibodies conjugated to HRP were detected using enhanced chemiluminescence (ECL, Amersham) with images captured on a luminescent image analyzer (LAS-3000, Fujifilm). Protein bands were quantified by densitometry using Image Studio™ Lite, v4.0 software.

### *In vivo* immunoprecipitation (IP)

FLAG-tagged human LRRK2 proteins were enriched from injected rat striatum using Protein G-Dynabeads (50 µl; Invitrogen) pre-coupled with mouse anti-FLAG-M2 antibody (5 µg; Sigma). Triton-soluble brain lysates (1.5 mg protein) were incubated with Dynabead-antibody complexes in 1 ml of IP buffer (1X PBS pH 7.4, 1% Triton X100) with rotation for 24 h at 4°C. Dynabead-antibody complexes were washed 3x with 1 ml IP buffer and 3x with 1X PBS and proteins were eluted at 70°C for 10 min in 2X Laemmli sample buffer (50 μl; Bio-Rad) containing 5% 2-mercaptoethanol. Immunoprecipitates were resolved by SDS-PAGE (4-20% Tris-Glycine gradient gels; Novex) and subjected to Western blotting with antibodies to pSer1292-LRRK2, pSer935-LRRK2, pSer910-LRRK2, total LRRK2 (clone N241A/34; NeuromAbs) and FLAG-M2 (Sigma).

### Antibodies

For Western blot and immunohistochemical analysis the following primary antibodies were used: mouse monoclonal anti-GFP (clones 7.1 and 13.1, Sigma), rabbit monoclonal anti-LRRK2 (clone N241A/34, NeuroMabs), rabbit monoclonal anti-LRRK2 (clone MJFF2/c41-2, abcam), mouse monoclonal anti-FLAG and anti-FLAG-HRP (clone M2, Sigma), rabbit monoclonal anti-pSer910-LRRK2 (clone UDD1 15(3), Abcam), rabbit monoclonal anti-pSer935-LRRK2 (clone UDD2 10(12), Abcam), rabbit monoclonal anti-pSer1292-LRRK2 (clone MJFR-19-7-8, Abcam), mouse monoclonal anti-amyloid precursor protein (APP, clone 22C11, Millipore), mouse monoclonal anti-α-Synuclein (Syn-1; clone 42, BD Biosciences), rabbit monoclonal anti-phospho-Ser129-α-Synuclein (clone EP1536Y, Abcam), rabbit polyclonal anti-Iba1 (019-19741, Wako), mouse monoclonal anti-β-tubulin (clone TUB 2.1, Sigma), rabbit polyclonal anti-dynamin-1 (PA1-660, ThermoFisher Scientific), rabbit polyclonal anti-tyrosine hydroxylase (NB300-109, Novus Biologicals), mouse monoclonal anti-ubiquitin (clone P4D1, Cell Signaling Technology), mouse monoclonal anti-phospho-Ser202/Thr205-tau (clone AT8; Pierce Biotech), mouse monoclonal anti-tau (clone TAU5, MAB361, Millipore), anti-rabbit polyclonal anti-glial fibrillary acidic protein (GFAP; E18320, Spring Bioscience), rabbit monoclonal anti-phospho-Thr73-RAB10 (MJF-R21, Abcam), rabbit monoclonal anti-RAB10 (D36C4, Cell Signaling Technology), mouse monoclonal anti-human adenovirus 5, hexon capsid protein (clone 9C12, Developmental Studies Hybridoma Bank), mouse monoclonal anti-GAPDH (clone 1E6D9, Protein Tech). For bright-field microscopy, biotinylated goat anti-mouse and anti-rabbit secondary antibodies (Vector Laboratories) were employed. For Western blots, light chain-specific mouse anti-rabbit IgG-HRP and goat anti-mouse IgG-HRP (Jackson Immunoresearch) secondary antibodies were used.

### Stereological analysis

The unbiased stereological estimation of substantia nigra dopaminergic neurons was performed using the optical fractionator probe of the StereoInvestigator software (MicroBrightField Biosciences), as described (32, 35, 44). For this analysis, every 4^th^ serial coronal section of 40 μm thickness, covering the whole region of the substantia nigra (typically from −4.5 mm to −6.6 mm relative to bregma) was immunostained with anti-TH antibody (NB300-109, Novus Biologicals) and counterstained with cresyl violet (Nissl) stain. Sampling was performed in a systematic random manner using a grid of 300 x 300 μm squares covering the substantia nigra pars compacta onto each section and applying an optical dissector consisting of a 65 x 65 x 16 μm square cuboid. All analyses were performed in a blind manner.

### Statistical analysis

One-way ANOVA with appropriate *post-hoc* analysis was used for multiple comparisons, whereas unpaired, two-tailed Student’s *t*-test was used for comparing two groups, as indicated. All data were plotted as mean ± SEM and were considered significant when *P*<0.05.

## Acknowledgements

This work was supported by funding from the National Institutes of Health (R01 NS091719; D.J.M.), Sanofi Aventis (D.J.M.), Michael J. Fox Foundation for Parkinson’s Research (D.J.M.), Swiss National Science Foundation (Grant No. 31003A_144063; D.J.M.) and American Parkinson Disease Association (A.P.T.N). The authors are grateful to Drs. Philippe Bertrand and Laurent Dubois (Sanofi, Chilly-Mazarin, France) for helpful discussions.

