## Supplemental Data for "Dopaminergic Neurodegeneration Induced by Parkinson’s Disease-Linked G2019S LRRK2 is Dependent on Kinase and GTPase Activity"

**Figure S1. Efficient retrograde transport of Ad5-GFP vector from striatum to nigral dopaminergic neurons in rat brain.** Ad5-eGFP vector was unilaterally delivered at six injection sites (1.5 x $10^{10}$ vp/site, in 2.5 µl) to the rat striatum. Immunofluorescence co-localization of GFP-positive neurons (anti-GFP, green) and TH-positive dopaminergic neurons (anti-TH, red) in the ipsilateral substantia nigra at 10 days post-injection. For the ipsilateral nigra, representative images from a rostro-caudal series are shown with each section separated by 80 µm from each other. Scale bars: 500 µm.

**Figure S2. Evaluation of human LRRK2 kinase activity following intrastriatal delivery of Ad5-LRRK2 vectors in rats. A)** Human LRRK2 levels and kinase activity in striatum. Ad5-LRRK2 (WT, G2019S or G2019S/K1906M) vectors were unilaterally delivered at six injection sites (1.5 x $10^{10}$ vp/site, in 2.5 µl) to the rat striatum. Ipsilateral striatum was harvested at 10 days post-delivery and subjected to sequential detergent extraction in 1% Triton X100 and 2% SDS. Striatal fractions were analyzed by Western blot analysis ($n$ = 3 animals/vector) indicating the levels of human LRRK2 variants (anti-FLAG antibody) or phosphorylated (Thr73) and total Rab10. GAPDH was used as a loading control. NI, non-injected striatum. Graphs indicate densitometric analysis of human LRRK2 variant levels (FLAG) normalized to GAPDH levels in each detergent fraction. Bars represent the mean ± SEM ($n$ = 3 animals/vector). *n.s*, non-significant by one-way ANOVA with Bonferroni's multiple comparisons. **B)** Ipsilateral striatal extracts from (A) were subjected to immunoprecipitation with anti-FLAG antibody to immunopurify human LRRK2 variants. Western blot analysis indicates levels of autophosphorylated (pSer1292), constitutively phosphorylated (pSer935) and total human LRRK2 (anti-FLAG antibody). Graphs indicate densitometric analysis of pSer1292 and pSer935 levels normalized to total FLAG-LRRK2 levels. Bars represent the mean ± SEM ($n$ = 3 animals/vector). *n.s*, non-significant by one-way ANOVA with Bonferroni's multiple comparisons. **C)** Ad5-LRRK2 (G2019S and G2019S/K1906M) vectors were unilaterally delivered at six injection sites (1.5 x $10^{10}$ vp/site, in 2.5 µl) to the rat striatum. Immunofluorescence co-localization of human LRRK2 (anti-FLAG antibody) and pSer1292-LRRK2 in ipsilateral striatum and nigra at 42 days post-injection. Arrowheads indicate FLAG-positive dopaminergic neurons (green), their corresponding pSer1292-LRRK2 signal (red), and merged images. Scale bars: 500 µm.

**Figure S3. Ad5-G2019S-LRRK2 induces axonal damage in the rat striatum. A-D)** Ad5-LRRK2 (WT, G2109S and G2019S/K1906M) vectors were unilaterally delivered at six injection sites (1.5 x $10^{10}$ vp/site, in 2.5 µl) to the rat striatum. **A)** Immunohistochemistry showing Ad5 capsid expression (anti-Hexon antibody) in the ipsilateral striatum at 42 days post-delivery. **B)** Immunohistochemistry showing increased ubiquitin-positive immunoreactivity (anti-ubiquitin antibody, clone P4D1) induced equivalently by all Ad5-LRRK2 vectors, and **C)** increased amyloid precursor protein-positive inclusions/spheroids (anti-

APP antibody, clone 22C11) induced by G2019S LRRK2 in ipsilateral striatum. **D)** Gallyas silver staining reveals increased axonal degeneration (black fibers) induced by G2019S LRRK2 in striosomal compartments of the ipsilateral striatum. Images indicate low and high magnification for each vector. Scale bars: 500 μm.

**Figure S4. Normal α-synuclein solubility, levels and Ser129 phosphorylation following intrastriatal delivery of Ad5-LRRK2 vectors in rats. A)** Western blot analysis of α-synuclein levels (Syn-1 antibody) or Ser129 phosphorylation (anti-pSer129-αSyn antibody) in 1% Triton-soluble or 2% SDS-soluble fractions from ipsilateral striatum and ventral midbrain of rats at 10 days after intrastriatal delivery of Ad5-LRRK2 vectors (WT, G2019S and G2019S/K1906M). Monomeric and high-molecular weight α-synuclein species are shown. GAPDH was used as a loading control. Molecular mass markers are indicated in kilodaltons. **B)** Densitometric analysis of pSer129-α-synuclein levels normalized to total α-synuclein, and total α-synuclein normalized to GAPDH for each detergent fraction. Bars represent the mean ± SEM (*n* = 3 animals/vector). **C)** Immunohistochemistry indicating a lack of Ser129 phosphorylated α-synuclein in substantia nigra at 42 days post-injection. Images indicate low and high magnification for each vector. Scale bars: 500 μm.

**Figure S5. LRRK2 variants do not alter tau solubility, levels or Ser202/Thr205 phosphorylation (AT8) following intrastriatal delivery of Ad5-LRRK2 vectors in rats. A)** Western blot analysis of tau levels (TAU5 antibody) or Ser202/Thr205 phosphorylation (AT8 antibody) in 1% Triton-soluble or 2% SDS-soluble fractions from ipsilateral striatum and ventral midbrain of rats at 10 days after intrastriatal delivery of Ad5-LRRK2 vectors (WT, G2019S and G2019S/K1906M). GAPDH was used as a loading control. Molecular mass markers are indicated in kilodaltons. **B)** Densitometric analysis of pSer202/Thr205-tau levels normalized to total tau, and total tau normalized to GAPDH for each detergent fraction. Bars represent the mean ± SEM (*n* = 3 animals/vector). **C)** Immunohistochemistry indicating increased Ser202/Thr205 phosphorylated tau (AT8-positive) in the ipsilateral substantia nigra at 42 days post-injection with each Ad5-LRRK2 vector. Images indicate low and high magnification for each vector. Scale bars: 500 μm.

**Figure S6. Ad5-LRRK2 vector delivery induces microgliosis in rat brain. A-B)** Ad5-LRRK2 (WT, G2019S and G2019S/K1906M) vectors were unilaterally delivered at six sites (1.5 x $10^{10}$ vp/site, in 2.5 μl) in the rat striatum. Immunohistochemistry indicates a modest increase in Iba1-positive microglial staining in the ipsilateral **A)** striatum and **B)** substantia nigra at 42 days post-injection. **C)** Ad5-LRRK2 (WT, G2019S and G2019S/K1906M) vectors were unilaterally delivered at one site (1.5 x $10^{10}$ vp/site, in 2.5 μl) in the rat substantia nigra. Immunohistochemistry reveals a marked increase of Iba1-positive microglia in the ipsilateral substantia nigra at 21 days post-injection. Images indicate low and high magnification for each vector. Scale bars: 500 μm.

**Figure S7. Endogenous LRRK2 is not required for efficient Ad5 vector infectivity in rodent brain.** Ad5-eGFP vectors were unilaterally delivered at four distinct sites (4.2 x $10^9$ vp/site, in 1 μl) to the striatum of adult wild-type (WT) and homozygous *LRRK2* knockout (KO) littermate mice. Immunohistochemistry indicates equivalent eGFP expression (anti-GFP antibody) in the ipsilateral striatum and substantia nigra (SNpc) of WT and KO mice at 10 days post-injection. Scale bars: 125 μm.

**Figure S8. Chronic in-diet dosing of PF-360 destabilizes human G2019S LRRK2 protein and inhibits endogenous LRRK2 kinase activity in rat brain. A)** Ad5-G2019S-LRRK2 vectors were unilaterally delivered at six distinct sites (1.5 x $10^{10}$ vp/site, in 2.5 μl) in the rat striatum. Rats were fed continuously with vehicle or PF-360 (175 mg/kg) chow from day 7 to 42 post-injection. Striatal tissues were harvested at 42 days post-injection and subjected to sequential extraction in 1% Triton X100 and 2% SDS. **B-C)** Western blot analysis of **B)** ipsilateral striatum extracts indicating human G2019S-LRRK2 levels (anti-FLAG antibody), or **C)** contralateral striatum extracts indicating phosphorylated (pSer935) and total (MJFF2/c41-2) endogenous LRRK2 levels in rats fed with vehicle or PF-360 chow. Dynamin I (DnmI) was used as a loading control for normalization. Densitometric quantitation is shown for **B)** human G2019S-LRRK2 (FLAG) levels in each detergent fraction, and **C)** endogenous pSer935-LRRK2 levels. Bars represent the mean ± SEM ($n$ = 4 animals/vector), *$P$<0.05 or **$P$<0.01 by unpaired Student *t*-test. **D)** Evaluation of body weight (grams) for individual rats (#1-12) monitored in 7-day periods fed with PF-360 (175 mg/kg) from day 7 to 42 post-injection. **E)** Total consumption of chow (grams) by individual rats (#1-12) monitored in 7-day periods fed with PF-360 (175 mg/kg) from days 7 to 42 post-injection.

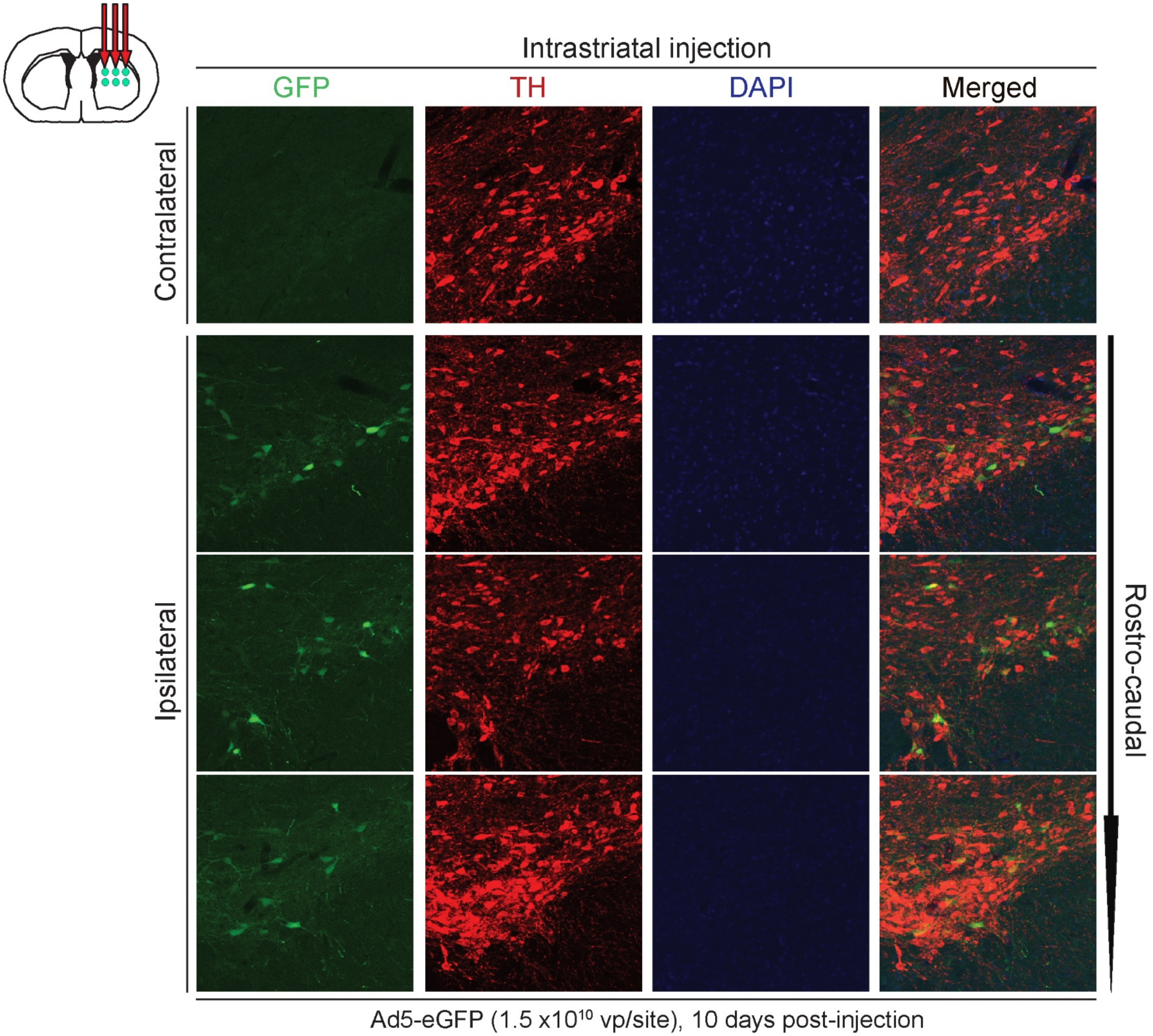
Intrastriatal injection
GFP
TH
DAPI
Merged
Contralateral
Ipsilateral
Rostro-caudal
Ad5-eGFP (1.5 x10^10 vp/site), 10 days post-injection

Figure S1

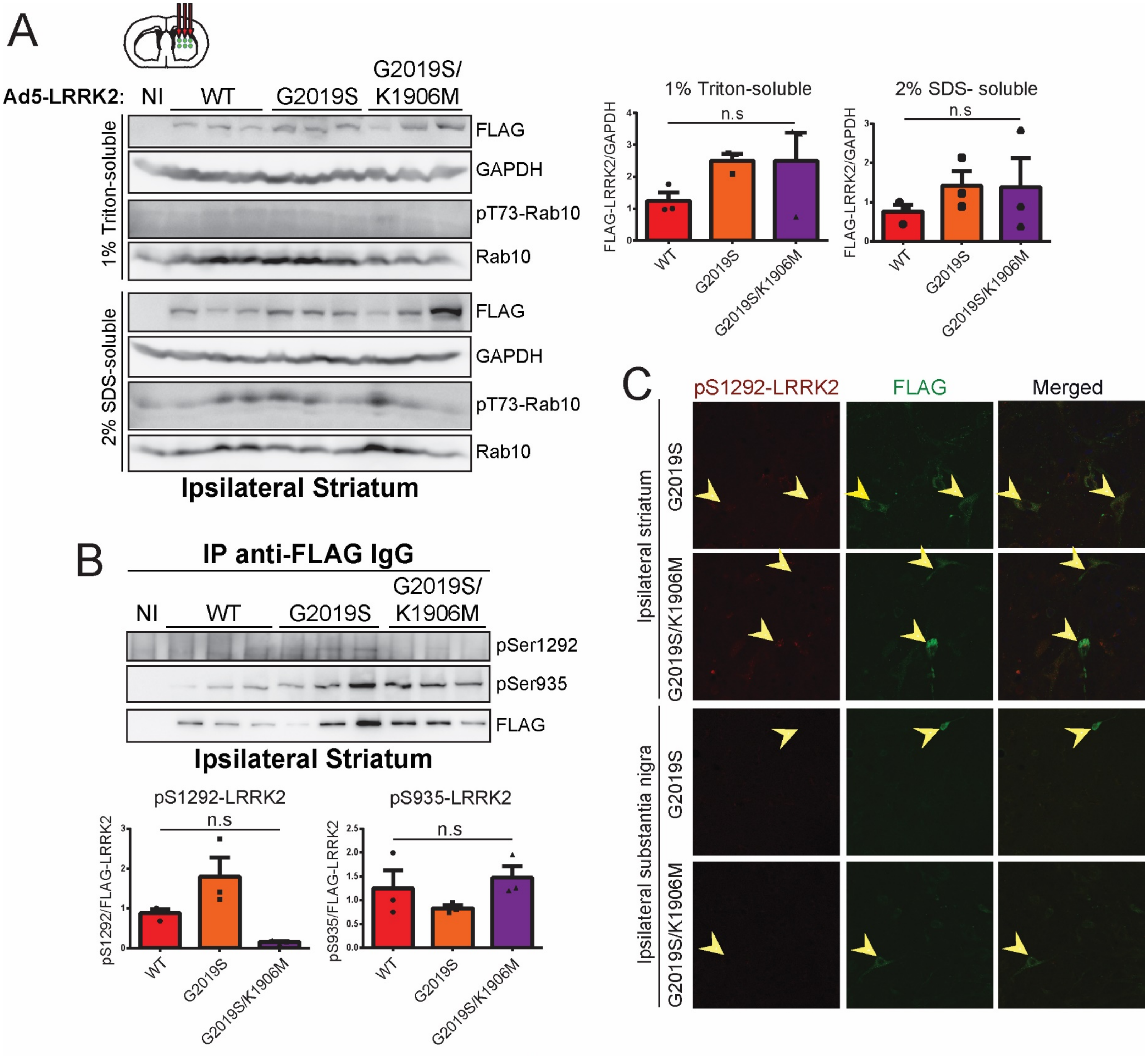
A
Ad5-LRRK2: NI WT G2019S G2019S/ K1906M
1% Triton-soluble
FLAG
GAPDH
pT73-Rab10
Rab10
2% SDS-soluble
FLAG
GAPDH
pT73-Rab10
Rab10
Ipsilateral Striatum
1% Triton-soluble
2% SDS- soluble
n.s
FLAG-LRRK2/GAPDH
WT
G2019S
G2019S/K1906M
B
IP anti-FLAG IgG
NI WT G2019S G2019S/ K1906M
pSer1292
pSer935
FLAG
Ipsilateral Striatum
pS1292-LRRK2
pS935-LRRK2
pS1292/FLAG-LRRK2
pS935/FLAG-LRRK2
C
pS1292-LRRK2
FLAG
Merged
Ipsilateral striatum
Ipsilateral substantia nigra
G2019S
G2019S/K1906M

Figure S2

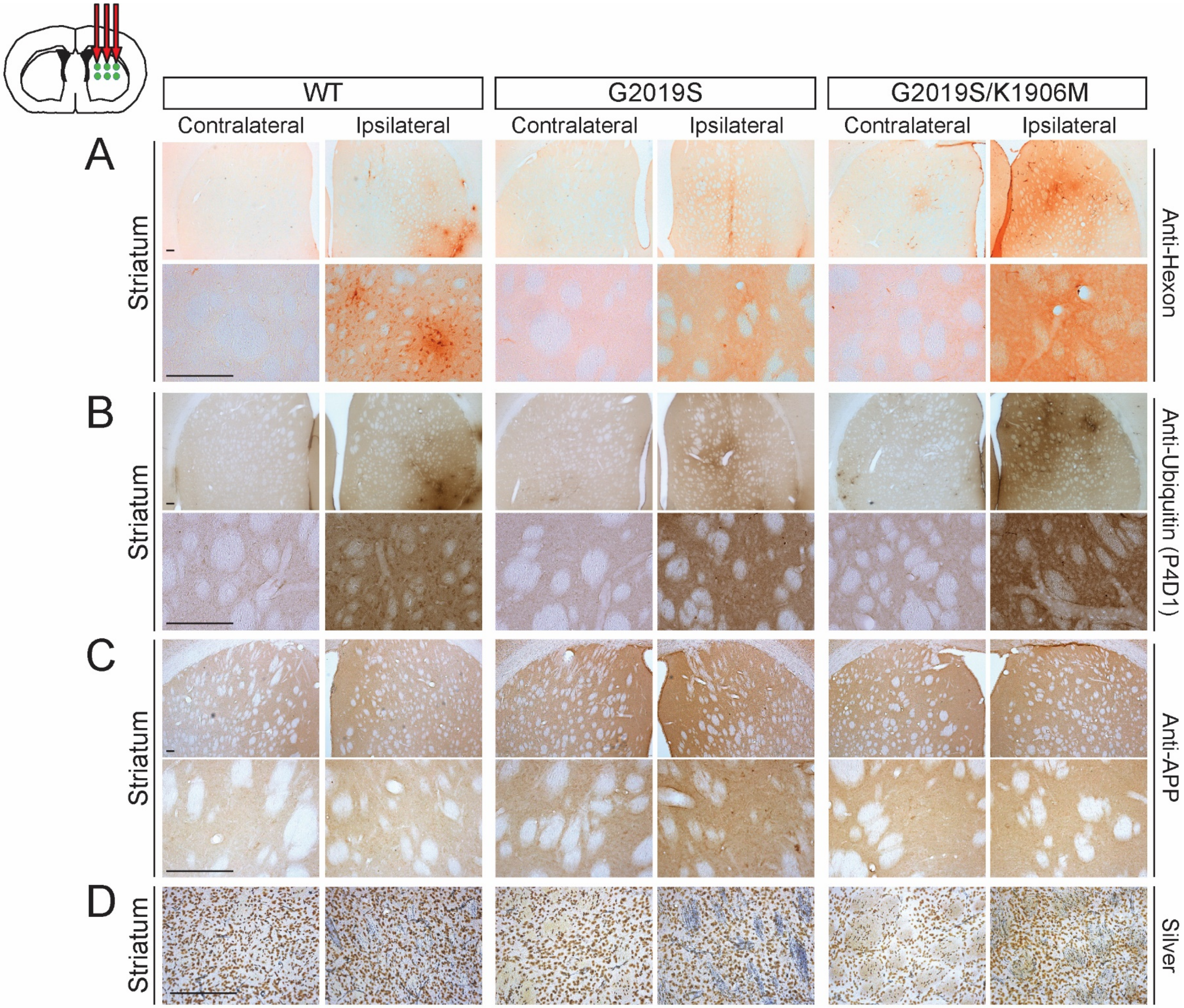
WT
G2019S
G2019S/K1906M
Contralateral
Ipsilateral
Contralateral
Ipsilateral
Contralateral
Ipsilateral
A
B
C
D
Striatum
Striatum
Striatum
Striatum
Anti-Hexon
Anti-Ubiquitin (P4D1)
Anti-APP
Silver

Figure S3

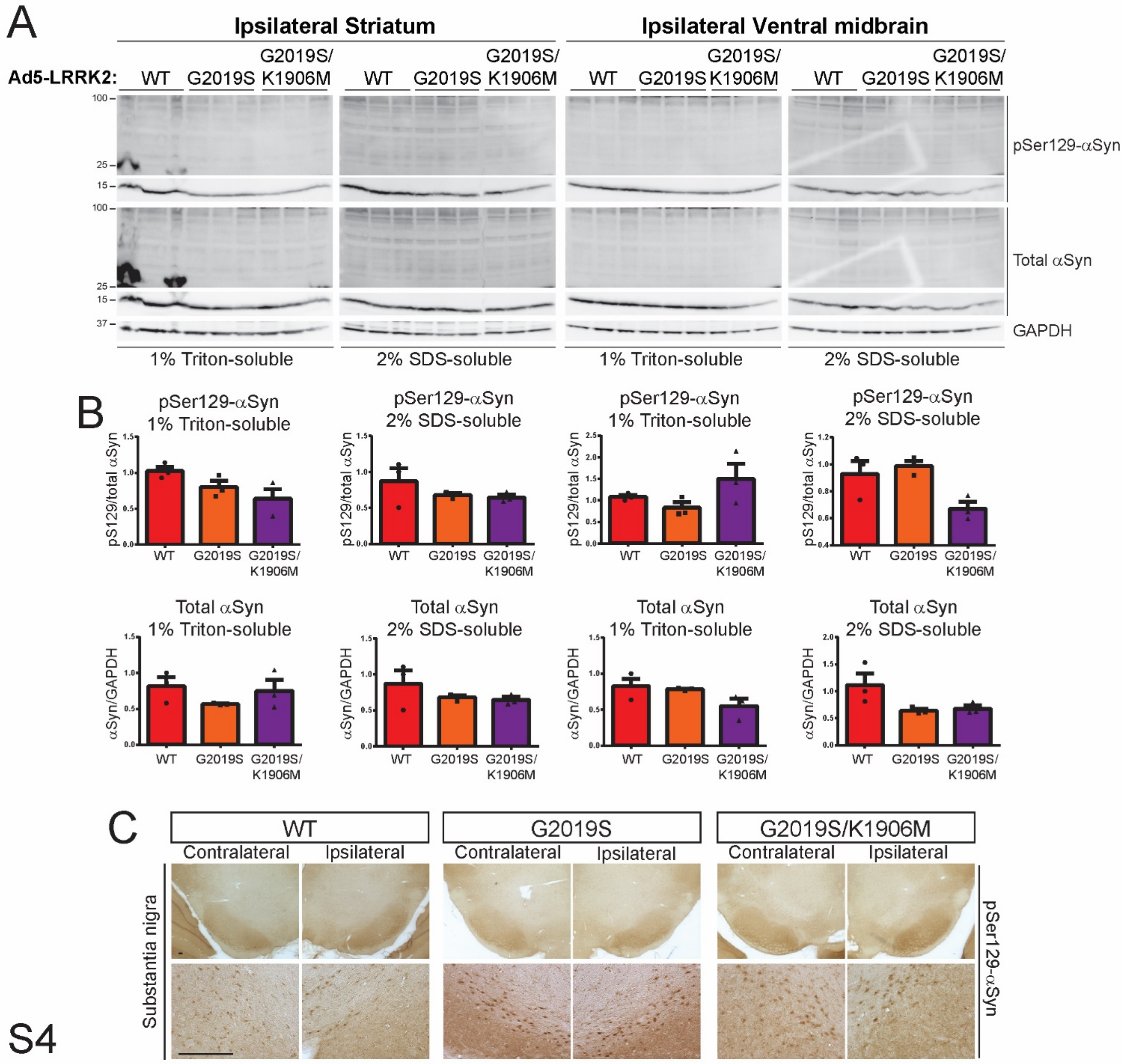
A
Ipsilateral Striatum
Ipsilateral Ventral midbrain
Ad5-LRRK2:
WT
G2019S
G2019S/ K1906M
pSer129-αSyn
Total αSyn
GAPDH
1% Triton-soluble
2% SDS-soluble
B
pSer129-αSyn 1% Triton-soluble
pSer129-αSyn 2% SDS-soluble
pS129/total αSyn
Total αSyn 1% Triton-soluble
Total αSyn 2% SDS-soluble
αSyn/GAPDH
C
Contralateral
Ipsilateral
Substantia nigra
G2019S/K1906M

Figure S4

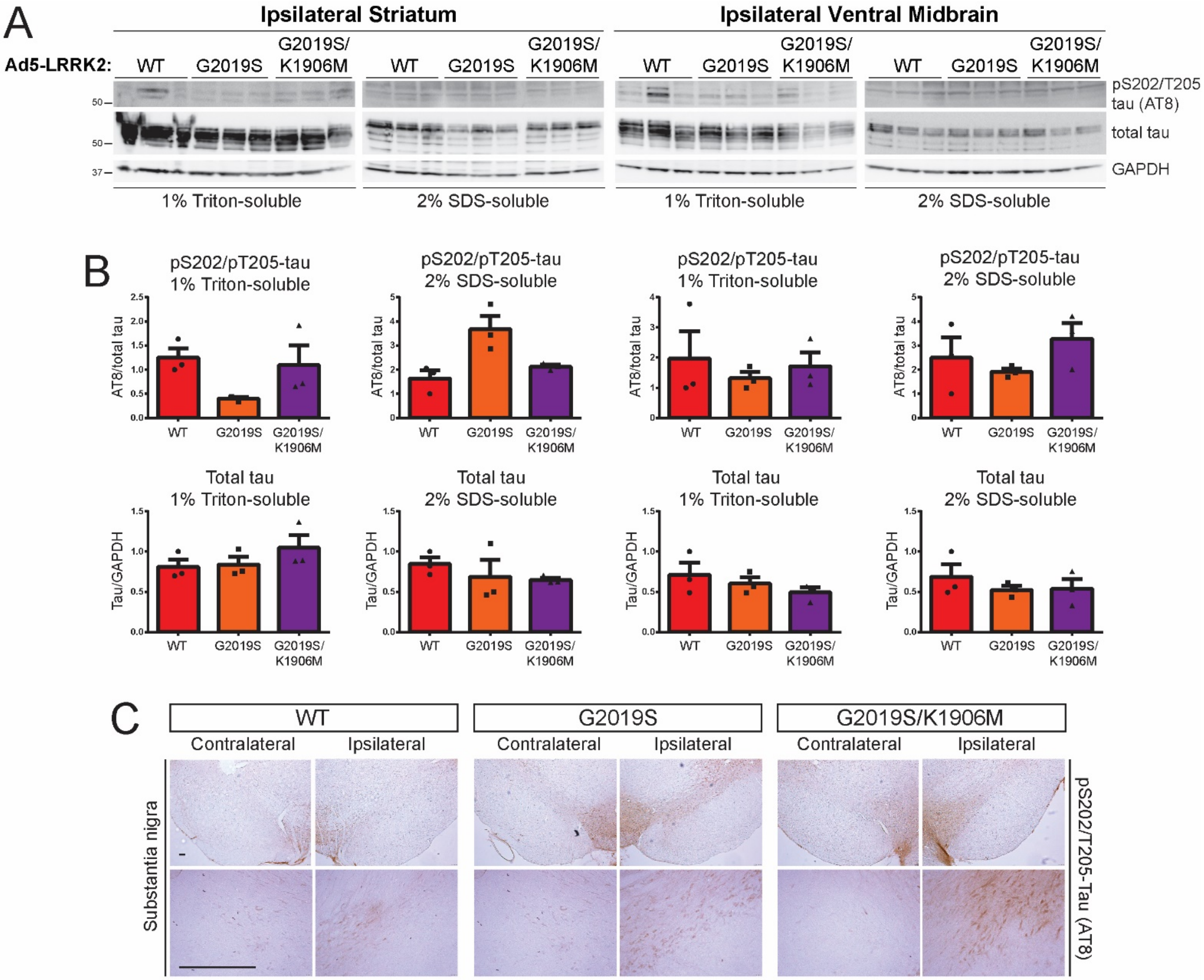
A
Ipsilateral Striatum
Ipsilateral Ventral Midbrain
Ad5-LRRK2:
WT
G2019S
G2019S/ K1906M
pS202/T205 tau (AT8)
total tau
GAPDH
50
50
37
1% Triton-soluble
2% SDS-soluble
B
pS202/pT205-tau 1% Triton-soluble
pS202/pT205-tau 2% SDS-soluble
AT8/total tau
Total tau 1% Triton-soluble
Total tau 2% SDS-soluble
Tau/GAPDH
C
Contralateral
Ipsilateral
G2019S/K1906M
Substantia nigra
pS202/T205-Tau (AT8)

Figure S5

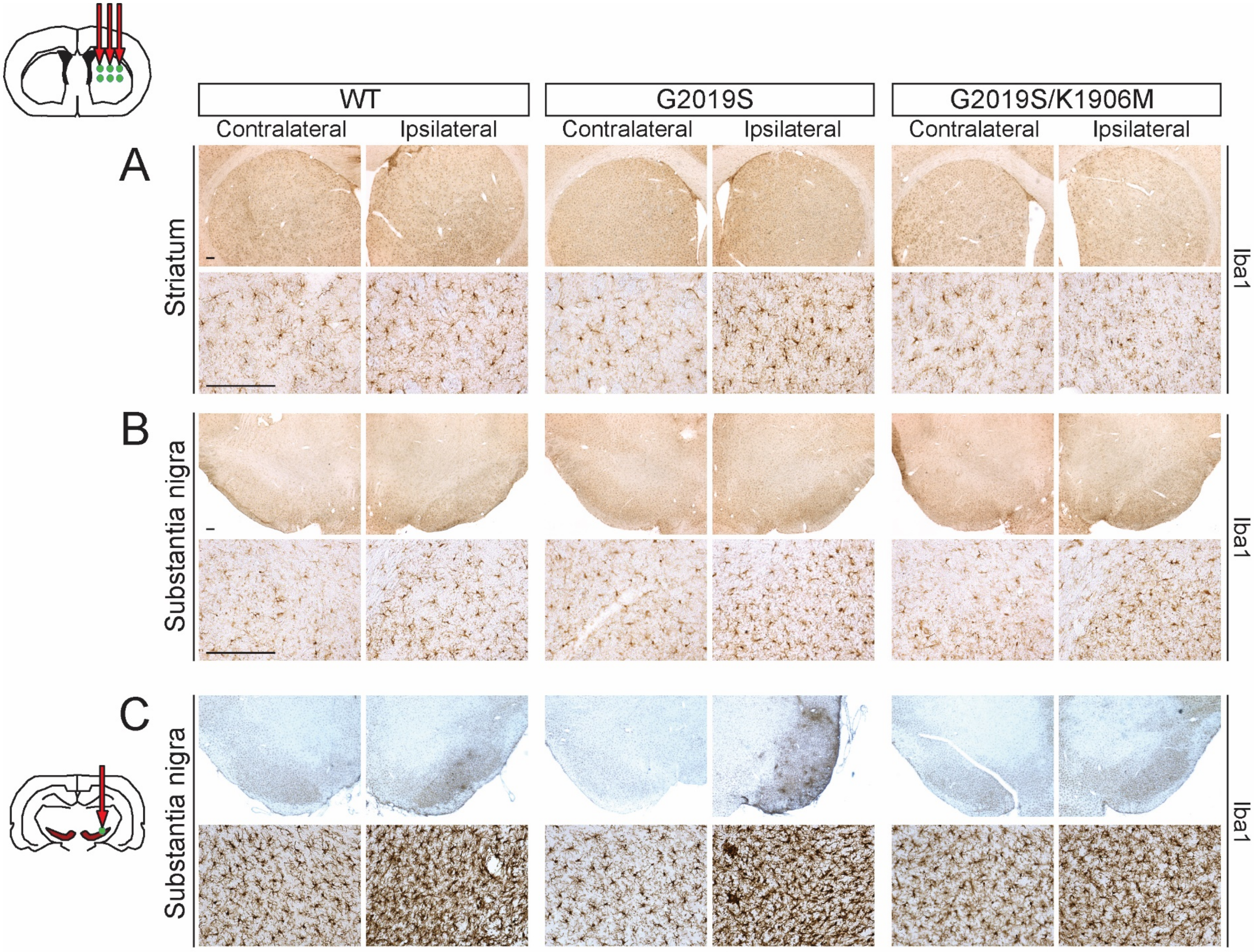
WT
G2019S
G2019S/K1906M
Contralateral
Ipsilateral
A
Striatum
B
Substantia nigra
C
Iba1

Figure S6

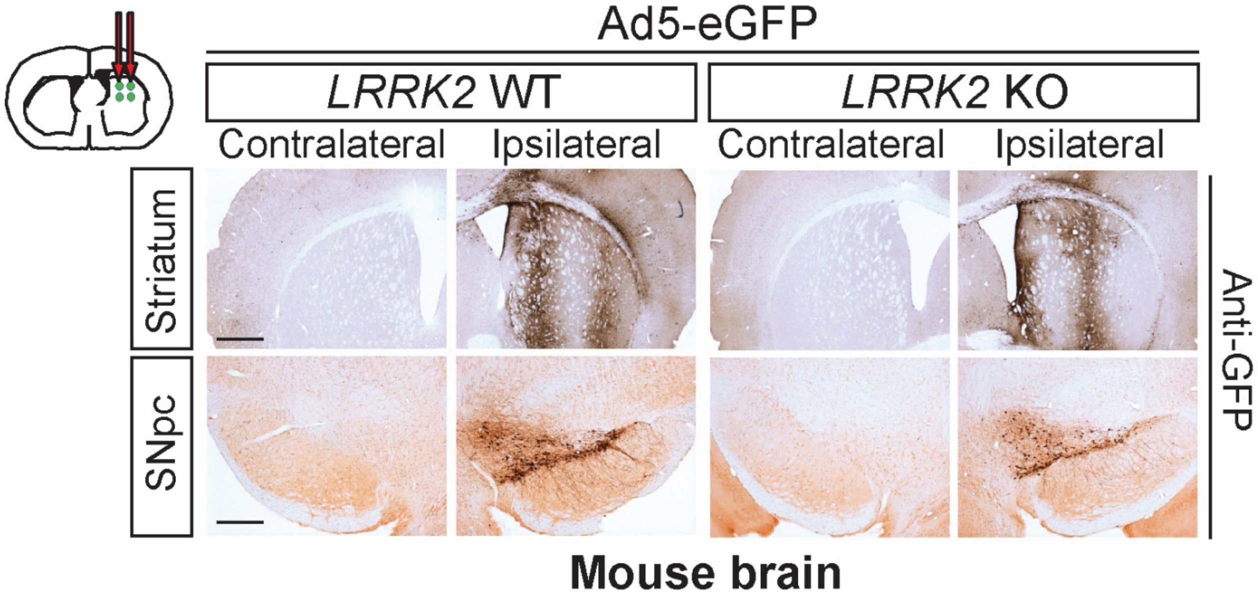
Ad5-eGFP
LRRK2 WT
LRRK2 KO
Contralateral
Ipsilateral
Contralateral
Ipsilateral
Striatum
SNpc
Anti-GFP
Mouse brain

Figure S7

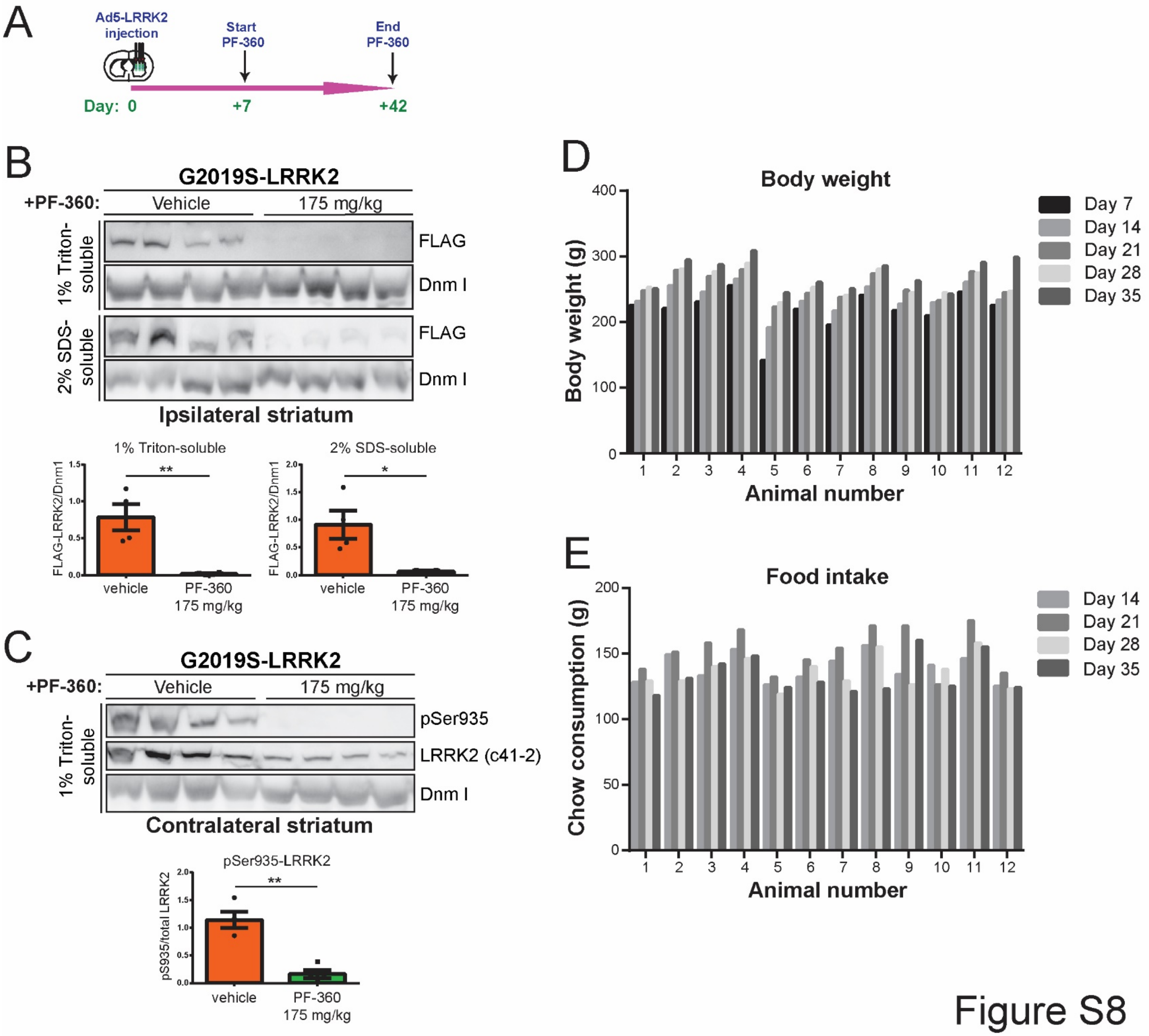
A
Ad5-LRRK2 injection
Start PF-360
End PF-360
Day: 0
+7
+42
B
G2019S-LRRK2
+PF-360:
Vehicle
175 mg/kg
1% Triton-soluble
2% SDS-soluble
FLAG
Dnm I
Ipsilateral striatum
1% Triton-soluble
2% SDS-soluble
FLAG-LRRK2/Dnm1
vehicle
PF-360 175 mg/kg
C
G2019S-LRRK2
+PF-360:
Vehicle
175 mg/kg
1% Triton-soluble
pSer935
LRRK2 (c41-2)
Dnm I
Contralateral striatum
pSer935-LRRK2
pS935/total LRRK2
vehicle
PF-360 175 mg/kg
D
Body weight
Body weight (g)
Day 7
Day 14
Day 21
Day 28
Day 35
Animal number
E
Food intake
Chow consumption (g)
Day 14
Day 21
Day 28
Day 35
Animal number

Figure S8
